# A high-performance genetically encoded fluorescent indicator for *in vivo* cAMP imaging

**DOI:** 10.1101/2022.02.27.482140

**Authors:** Liang Wang, Chunling Wu, Wanling Peng, Ziliang Zhou, Jianzhi Zeng, Xuelin Li, Yini Yang, Shuguang Yu, Ye Zou, Mian Huang, Chang Liu, Yefei Chen, Yi Li, Panpan Ti, Wenfeng Liu, Yufeng Gao, Wei Zheng, Shangbang Gao, Zhonghua Lu, Pei-Gen Ren, Ho Leung Ng, Jie He, Shoudeng Chen, Min Xu, Yulong Li, Jun Chu

## Abstract

cAMP is a key second messenger that regulates diverse cellular functions including neural plasticity. However, the spatiotemporal dynamics of intracellular cAMP in intact organisms are largely unknown due to low sensitivity and/or brightness of current genetically encoded fluorescent cAMP indicators. Here, we report the development of the new circularly permuted GFP (cpGFP)-based cAMP indicator G-Flamp1, which exhibits a large fluorescence increase (a maximum ΔF/F_0_ of 1100% in HEK293T cells), relatively high brightness, appropriate affinity (a K_d_ of 2.17 µM) and fast response kinetics (an association and dissociation half-time of 0.20 s and 0.087 s, respectively). Furthermore, the crystal structure of the cAMP-bound G-Flamp1 reveals one linker connecting the cAMP-binding domain to cpGFP adopts a distorted β-strand conformation that may serve as a fluorescence modulation switch. We demonstrate that G-Flamp1 enables sensitive monitoring of endogenous cAMP signals in brain regions that are implicated in learning and motor control in living organisms such as fruit flies and mice.

## Introduction

Cyclic adenosine 3’,5’-monophosphate (cAMP), which is produced from adenosine triphosphate (ATP) by adenylyl cyclase (AC), acts as a key second messenger downstream of many cell surface receptors, especially G-protein-coupled receptors (GPCRs)^1^. cAMP plays critical roles in regulating numerous cellular physiological processes, including neuronal plasticity and innate and adaptive immune cell activities, through its effector proteins such as protein kinase A (PKA), exchange protein directly activated by cAMP (EPAC) and cyclic nucleotide-activated ion channels (CNG and HCN channels)^2^. A growing body of evidence has shown that cAMP is precisely controlled in space and time in living cells and its abnormal dynamics are associated with many diseases^3^. However, it is largely unclear how cAMP signaling is regulated under physiological and pathological conditions *in vivo*^3–5^.

Genetically encoded fluorescent indicators (GEFIs) with advanced optical imaging have emerged as a powerful tool for real-time monitoring the spatiotemporal dynamics of signaling molecules including calcium in intact model organisms^6^. Current GEFIs for cAMP were developed based on two strategies: fluorescence resonance energy transfer (FRET) between two fluorescent proteins (FPs) or circular permutation/splitting of a single FP^7–9^. The latter is much more sensitive and, because they only require a single-color channel, can be more easily used together with other spectrally compatible sensors and actuators^10^. So far, a few single-FP cAMP sensors (Flamindo2, cAMPr, Pink Flamindo and R-FlincA) based on different mammalian cyclic nucleotide-binding domains (CNBDs) and green/red FPs have been created^11–14^. However, they exhibit small fluorescence changes (|ΔF/F_0_| < 150%) and most are dim in mammalian cells at 37 °C (Supplementary Fig. 1a-c). Thus, it is highly desirable to develop new high-performance (high brightness, high sensitivity and fast response kinetics) single-FP cAMP sensors that can decipher complex cAMP signals *in vivo*.

To address these problems, we engineered a highly responsive circularly permuted GFP (cpGFP)-based cAMP sensor named G-Flamp1 (<u>g</u>reen <u>fl</u>uorescent c<u>AMP</u> indicator <u>1</u>) by inserting cpGFP into the CNBD of the bacterial *Mloti*K1 channel (mlCNBD), followed by extensive screening. G-Flamp1 exhibits a maximum ΔF/F_0_ of 1100% in HEK293T cells at 37 °C, which is 9−47 times greater than existing single-FP cAMP sensors. Furthermore, we resolved the crystal structure of cAMP-bound G-Flamp1 and found a long distorted β-strand connecting mlCNBD and cpGFP, which is unseen in other single-FP sensors and could critically modulate sensor fluorescence. Finally, we successfully monitored cAMP signals with G-Flamp1 during learning and motor control in fruit flies and mice.

## Results

### Development of G-Flamp1

To develop a high-performance genetically encoded cAMP indicator (GEAI), we chose mlCNBD as a starting point (Fig. 1a)^15, 16^. Unlike mammalian CNBDs, the bacterial mlCNBD likely does not interact with endogenous eukaryotic proteins and thus would not interfere with signaling pathways in mammalian cells^8^. Furthermore, mlCNBD exhibits high binding affinity and specificity for cAMP because the dissociation constants (K_d_) for mlCNBD-cAMP and mlCNBD-cGMP complexes are 68 nM and 499 nM, respectively. In addition, it has fast response kinetics with an association half-time (t_on_) of 27 ms under 1 µM cAMP and dissociation half-time (t_off_) of 74 ms^16^. Lastly, the tertiary structure of mlCNBD, especially the cAMP-binding pocket, is significantly different from those of mammalian CNBDs (Supplementary Fig. 2), raising the possibility that cAMP sensors with a different response profile can be engineered.

**Fig. 1.**
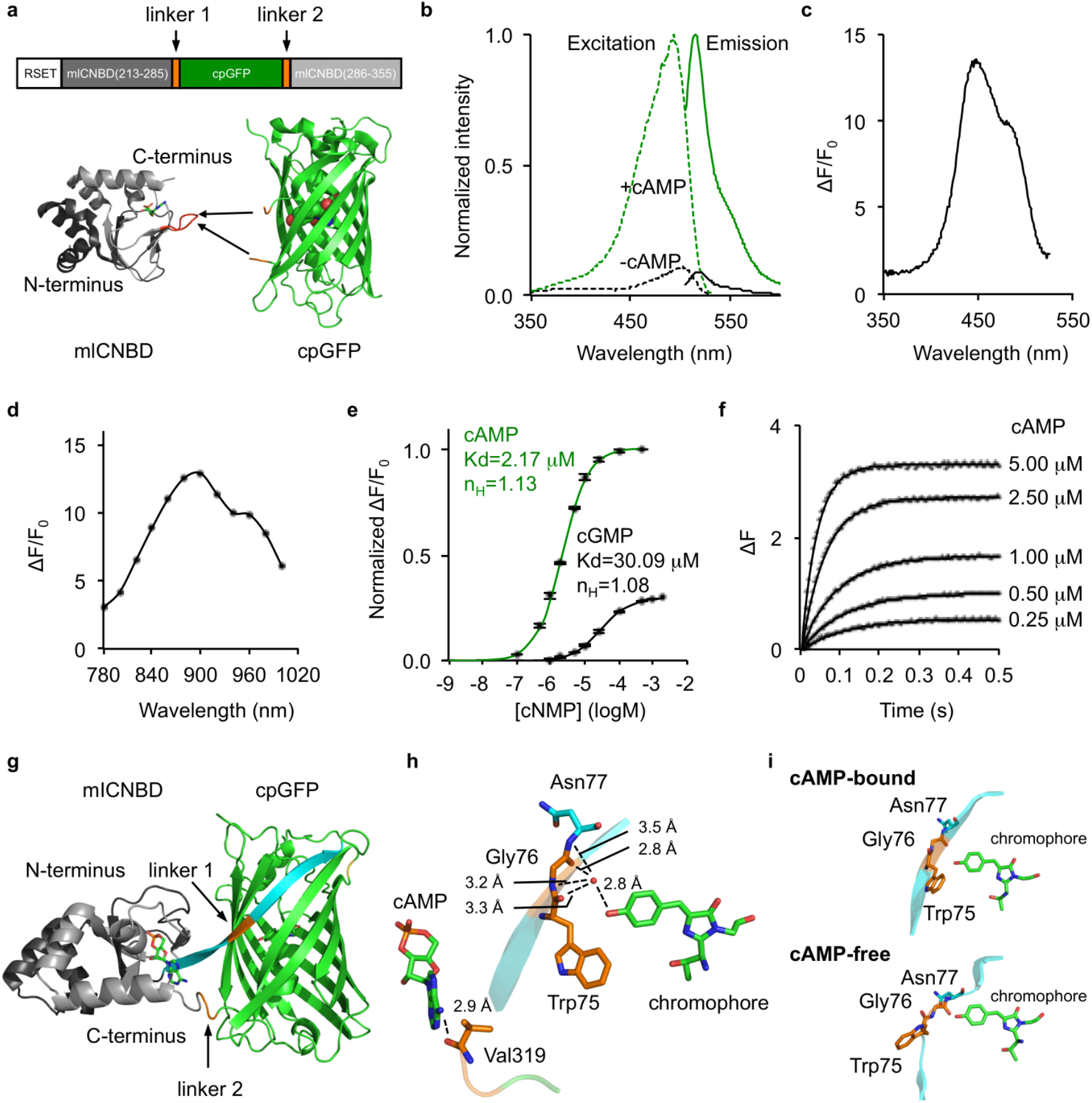
Development and *in vitro* characterization of G-Flamp1 indicator. (**a**) Schematic of G-Flamp sensors. cpGFP with two flanking linkers (two amino acids per linker) is inserted into mlCNBD (Gly213-Ala355, Genbank accession number: BA000012.4). The N-terminal peptide (RSET) including a 6× His tag is from the bacterial expression vector pNCS. The X-ray crystal structures of cAMP-bound mlCNBD (PDB: 1VP6) and cpGFP (PDB: 3WLD) are shown as cartoon with cAMP and chromophore of cpGFP shown as stick and sphere, respectively. The loop bearing the insertion site in G-Flamp1 is marked in red. (**b**) Excitation and emission spectra of cAMP-free and cAMP-bound G-Flamp1 sensors in HEPES buffer (pH 7.15). (**c**) Excitation wavelength-dependent ΔF/F_0_ of G-Flamp1 under one-photon excitation. (**d**) Excitation wavelength-dependent ΔF/F_0_ of G-Flamp1 under two-photon excitation. (**e**) Binding titration curves of G-Flamp1 to cAMP or cGMP in HEPES buffer (pH 7.15). The data were fitted by a sigmoidal binding function to extract the dissociation constant K_d_ and Hill coefficient n_H_. Data are presented as mean ± standard error of mean (SEM) from three independent experiments. (**f**) Binding kinetics of G-Flamp1 to cAMP measured using the stopped-flow technique in HEPES buffer (pH 7.15). Each curve corresponds to a different concentration of cAMP, i.e., from bottom to top: 0.25 μM, 0.5 μM, 1 μM, 2.5 μM and 5 μM. The data were fitted by a single-exponential function. (**g**) Cartoon representation of crystal structure of cAMP-bound G-Flamp1 (PDB: 6M63). The N-and C-terminal fragments of mlCNBD are shown in dark and light grey, respectively. cpGFP is in green and both linkers are in orange. The long β-strand possessing linker 1 is in cyan. (**h**) Chromophore and cAMP are in close proximity with linker 1 and linker 2, respectively. (**i**) Zoom-in view of linker 1 and its neighboring residues and the chromophore in the cAMP-bound crystal structure and the simulated cAMP-free structure.

To determine the optimal insertion site, we varied the position of cpGFP with original linkers from the calcium sensor GCaMP6f (LE-cpGFP-LP)^17^ in three loop regions of mlCNBD: the region ‘Gln237-Leu239’ undergoes a large conformation change from random coil to α-helix upon cAMP binding while the regions ‘Ala283-Val288’ and ‘Ala313-Val317’ remain random coils with small conformation change (numbering according to PDB 1VP6 of mlCNBD). A total of 11 sensors were tested using a bacterial lysate screening assay. One dim variant named G-Flamp0.1, in which cpGFP was inserted between Pro285 and Asn286 of mlCNBD, gave the largest signal change with a ΔF/F_0_ of −25.8% (Fig. 1a and Supplementary Fig. 3a-c). To improve the brightness of G-Flamp0.1, we examined several beneficial mutations from the well-folded GFP variants (Citrine and superfolder GFP)^18, 19^ and generated G-Flamp0.2 with brightness increased by 330% (Supplementary Fig. 3d). To obtain a large ΔF/F_0_ sensor, both linkers connecting cpGFP and mlCNBD were randomized together. Of the 427 variants tested, one variant (G-Flamp0.5) with linkers ‘WG’ and ‘RV’ showed the largest fluorescence change with a ΔF/F_0_ of 230% when excited at 488 nm (Supplementary Fig. 3e). Next, we performed random mutagenesis on G-Flamp0.5 using error-prone PCR and were able to identify a bright and highly responsive variant G-Flamp0.7 with a ΔF/F_0_ of 560%, which harbors P285N mutation in mlCNBD and D173G mutation in GFP (Supplementary Fig. 3f-g). Finally, to increase the selectivity for cAMP over cGMP (defined as K_d_ ratio of cGMP/cAMP), the mutation S308V, which is in the cAMP-binding pocket, was introduced to weaken the binding between mlCNBD and cGMP^20^. The resultant sensor G-Flamp1 had a higher selectivity with a ΔF/F_0_ of 820% under excitation at 488 nm (Supplementary Fig. 3g and Supplementary Fig. 4).

### *In vitro* characterization of G-Flamp1 sensor

We first investigated the fluorescence and absorption properties of purified G-Flamp1. The cAMP-bound G-Flamp1 had excitation and emission peaks at 490 nm and 510 nm, respectively, which were similar to those of mEGFP. The excitation and emission peaks of cAMP-free G-Flamp1 were redder than those of cAMP-bound G-Flamp1 by 10 nm and 3 nm, respectively (Fig. 1b and Supplementary Fig. 5a-b), suggesting different chromophore environments in cAMP-bound and cAMP-free G-Flamp1. According to these fluorescence spectra, the calculated fluorescence change peaked at 450 nm with a maximum ΔF/F_0_ of 1300% (Fig. 1c). Absorbance spectra revealed that both cAMP-bound and cAMP-free G-Flamp1 displayed two peaks with maxima at 400 nm and 490 nm (cAMP-bound G-Flamp1) or 500 nm (cAMP-free G-Flamp1) (Supplementary Fig. 5c), which correspond to protonated (dark state) and deprotonated (bright state) chromophores, respectively^21^. Moreover, the deprotonated form of cAMP-bound G-Flamp1 significantly increased, making it much brighter than deprotonated cAMP-free G-Flamp1. Under two-photon illumination, cAMP-bound G-Flamp1 had a similar excitation spectrum to mEGFP with the peak at around 920 nm (Supplementary Fig. 5d) and a maximum ΔF/F_0_ of 1300% at around 900 nm (Fig. 1d).

Compared to cAMP-free G-Flamp1, cAMP-bound G-Flamp1 exhibited a 6-fold greater extinction coefficient (EC) (25280 mM^-1^cm^-1^ versus 4374 mM^-1^cm^-1^) and similar quantum yield (QY) (0.322 versus 0.323) (**Supplementary Table 1**). Like other single-FP probes, the fluorescence intensity of G-Flamp1 was sensitive to pH, with pKa values of 8.27 and 6.95 for cAMP-free and cAMP-bound G-Flamp1, respectively (Supplementary Fig. 6a). Moreover, the calculated ΔF/F_0_ peaked at pH 6.5 with a value of 1640% and remained high at pH 7.0 with a value of 1440% (Supplementary Fig. 6b), indicating that G-Flamp1 would be highly responsive in mammalian cells where the physiological pH is maintained between 6.8−7.3^22^.

The dose-response curves showed that K_d_ values of G-Flamp1 for cAMP and cGMP were 2.17 µM and 30.09 µM, respectively (Fig. 1e), leading to a 13-fold higher selectivity for cAMP over cGMP, which is similar to other widely used cAMP probes (**Supplementary Table 2**)^14^. Since the K_d_ value for G-Flamp1-cAMP complex is close to the resting cAMP concentration of 0.1−1 µM^23, 24^, G-Flamp1 should detect cAMP changes under physiological stimulation conditions. To measure response kinetics, we applied the stopped-flow technique on purified G-Flamp1 and fitted data with a mono-exponential function. The apparent association (k_on_) and dissociation (k_off_) rate constants were 3.48 µM^-1^s^-1^ and 7.9 s^-1^, resulting in a t_on_ of 0.20 s under 1 µM cAMP and t_off_ of 0.087 s, respectively (Fig. 1f). Taken together, these results indicate that G-Flamp1 can faithfully report cAMP dynamics with sub-second temporal resolution.

### Crystal structure of cAMP-bound G-Flamp1

To understand the molecular mechanism of large fluorescence change in G-Flamp1 indicator, we determined the X-ray crystal structure of cAMP-bound G-Flamp1 without RSET tag at pH 8.0 to a 2.2 Å resolution (Fig. 1g). The statistics of data collection and structure refinement were summarized in **Supplementary Table 3**. Overall, all residues in G-Flamp1 showed good electron density except for N-terminal nine residues (MGFYQEVRR), C-terminal six residues (GAAASA) and a flexible linker (GGTGGS) within cpGFP. Two G-Flamp1 molecules were arranged as a dimer in one asymmetric unit of G-Flamp1 crystal and were structurally similar with an r.m.s.d. of Cα atoms of 0.149 Å. However, this homodimer was not biologically relevant and is likely caused by crystallographic packing because its dimerization interface is mediated by β-barrel ends of cpGFP rather than the previously described β-barrel wall^25^.

The linkers connecting sensing domain and circularly permuted FP (cpFP) are the main determinant of the dynamic range of single-FP sensors^9^. The crystal structure of cAMP-bound G-Flamp1 reveals that the first linker Trp75/Gly76 and the second linker Arg318/Val319 (numbering according to PDB 6M63 of G-Flamp1, Supplementary Fig. 4c), along with their flanking amino acids from mlCNBD and cpGFP, adopt a highly twisted β-strand and random coil conformation, respectively (Fig. 1g and Supplementary Fig. 7), which is unique because both linkers in other single-FP sensors with crystal structures available fold as random coil or α-helix segments (Supplementary Fig. 8)^26–28^. In G-Flamp1, linker 1 and linker 2 are in close proximity with chromophore and cAMP, respectively (Fig. 1h), suggesting the former primarily contributes to fluorescence change. Moreover, since the mlCNBD domain is far away from the chromophore, we reasoned that a self-contained fluorescence modulation mechanism, in which residues from linkers and/or FP (e.g., the red calcium sensor K-GECO1) rather than sensing domain (e.g., the green calcium sensor GCaMP3) interact with the deprotonated chromophore (Supplementary Fig. 8)^26^, may exist in G-Flamp1.

A close examination of linker 1 revealed that the Trp75 stabilizes the phenolic group of the chromophore in two ways. First, the backbone CO or NH groups of the tripeptide Trp75-Gly76-Asn77 indirectly interact with the phenolic oxygen of the chromophore, via a water molecule, to form a hydrogen-bonding network. Second, the bulky side chain of Trp75 protects the chromophore from solvent. Thus, we reasoned that a movement of Trp75 would make the chromophore unstable and dim. Consistent with this, molecular dynamics (MD) simulations of cAMP-free G-Flamp1 showed that the tripeptide underwent significant conformational rearrangement with a transition from β-strand to random coil and the side chain of Trp75 rotating away from the chromophore while linker 2 showed small change (Fig. 1i and **Supplementary Video 1**). Subsequent saturation mutagenesis on position 75 demonstrated that all G-Flamp1 variants had reduced fluorescence changes with a ΔF/F_0_ of 0%−232% (Supplementary Fig. 9), further confirming the critical role of Trp75 in tuning fluorescence change of G-Flamp1 in a self-contained manner. However, to verify these assumptions, a crystal structure of cAMP-free G-Flamp1 needs to be resolved and compared to that of cAMP-bound G-Flamp1.

### Performance of G-Flamp1 in mammalian cells

We first examined the cellular localization and brightness of G-Flamp1 in HEK293T cells. G-Flamp1, like Flamindo2 and Pink Flamindo, was evenly distributed in cytoplasm and nucleus (Fig. 2a and Supplementary Fig. 1b. The detailed imaging conditions throughout the paper are summarized in **Supplementary Table 4**). In contrast, cAMPr and R-FlincA were found to localize mainly in the cytosol (Fig. 2a and Supplementary Fig. 1b), with the latter forming puncta 48 hours post transfection (Supplementary Fig. 1d) and thus likely being toxic to mammalian cells^26^. Under one-photon (488 nm) illumination, the basal fluorescence intensities of G-Flamp1, cAMPr and Flamindo2 were 49%, 109% and 11% of that of GCaMP6s^17^, respectively (Fig. 2a-b). At 450 nm, which gives the largest ΔF/F_0_, the brightness is reduced by half and is ∼25% of that of GCaMP6s taking the excitation efficiencies at 450 nm and 488 nm into account (Fig. 1b). Again, under two-photon (920 nm) illumination, G-Flamp1 was brighter than Flamindo2 but dimmer than cAMPr (74% versus 38% versus 165% of GCaMP6s) in the resting state (Supplementary Fig. 10a).

**Fig. 2.**
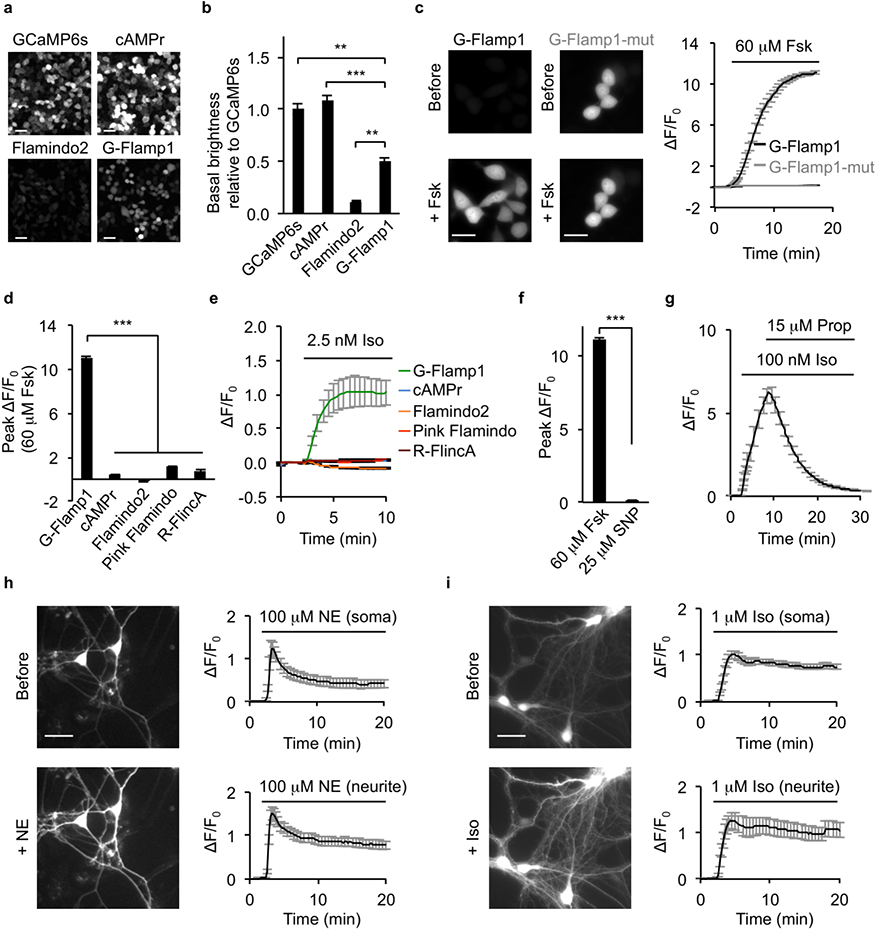
Characterization of G-Flamp1 in mammalian cells. (**a**) Wide-field fluorescence images of green cAMP sensors (cAMPr, Flamindo2 and G-Flamp1) and GCaMP6s in resting HEK293T cells. Scale bars: 50 µm. (**b**) Brightness of GCaMP6s, cAMPr, Flamindo2 and G-Flamp1 in resting HEK293T cells measured using a plate reader. n = 3 wells of the 24-well plates for each indicator. Two-tailed Student’s *t*-tests were performed. *P* = 0.0018, 2.6 × 10^-4^ and 0.0036 between G-Flamp1 and GCaMP6s, cAMPr and Flamindo2, respectively. (**c**) Representative fluorescence images (left) and traces of ΔF/F_0_ (right) in response to 60 µM Fsk in HEK293T cells expressing G-Flamp1 or G-Flamp1-mut. n = 34 cells (G-Flamp1) and 22 cells (G-Flamp1-mut) from 3 independent experiments. Scale bars: 20 µm. (**d**) Peak ΔF/F_0_ in response to 60 µM Fsk in HEK293T cells expressing G-Flamp1 or other cAMP sensors. n = 34 cells (G-Flamp1), 33 cells (cAMPr), 42 cells (Flamindo2), 34 cells (Pink Flamindo) and 18 cells (R-FlincA) from 3 independent experiments. Two-tailed Student’s *t*-tests were performed. *P* = 3.5 × 10^-38^, 2.8 × 10^-38^, 1.4 × 10^-41^ and 1.6 × 10^-46^ between G-Flamp1 and cAMPr, Flamindo2, Pink Flamindo and R-FlincA, respectively. (**e**) Representative traces of ΔF/F_0_ in response to 2.5 nM Iso in HEK293T cells expressing different cAMP sensors. n = 30 cells (G-Flamp1), 28 cells (cAMPr), 27 cells (Flamindo2), 27 cells (Pink Flamindo) and 14 cells (R-FlincA) from 3 independent experiments. (**f**) Peak ΔF/F_0_ in response to 60 µM Fsk or 25 µM SNP in HEK293T cells expressing G-Flamp1. n = 34 cells (Fsk) and 15 cells (SNP) from 3 independent experiments. Two-tailed Student’s *t*-test was performed. *P* = 4.3 × 10^-38^ between Fsk and SNP treatments. (**g**) Representative traces of ΔF/F_0_ in response to 100 nM Iso followed by 15 µM propranolol in HEK293T cells expressing G-Flamp1. n = 17 cells from 3 cultures. (**h-i**) Representative fluorescence images (left) and traces of ΔF/F_0_ in response to 100 µM NE (h) or 1 µM Iso (i) in cortical neurons expressing G-Flamp1. n = 10 (soma) and 9 (neurite) regions of interest (ROIs) of 10 neurons from 3 cultures in **h** and n = 28 (soma) and 14 (neurite) ROIs of 28 neurons from 3 cultures in **i**. Scale bars: 20 µm. Data are presented as mean ± SEM in **b**, **c** (right), **d**, **e**, **f**, **g**, **h** (right) and **i** (right). \*\*\**P* < 0.001 and \*\**P* < 0.05.

Next we evaluated the cytotoxicity and interference with cAMP signaling of G-Flamp1 at a medium expression level. HEK293T cells stably expressing G-Flamp1 proliferated similarly to untransfected cells (Supplementary Fig. 11a), suggesting low cytotoxicity of G-Flamp1. To assess G-Flamp1’s buffering effect, we investigated the phosphorylation of cAMP response element binding protein (CREB) at Ser133, a key molecular event downstream of cAMP-PKA^29^. Both G-Flamp1-expressing HEK293T and control cells showed similar basal levels and increases of phospho-S133 of CREB before and after 10 µM β-adrenergic receptor (β-AR) agonist isoproterenol (Iso) stimulation, respectively (Supplementary Fig. 11b). Taken together, these results indicate that G-Flamp1 expression had no obvious effects on endogenous signaling.

We further determined the fluorescence change and sensitivity of G-Flamp1. Forskolin (Fsk), a potent activator of transmembrane AC^30^, was used to induce a high level of cAMP to assess the maximum fluorescence change. Under 450 nm illumination, G-Flamp1 expressed in HEK293T cells exhibited a maximum ΔF/F_0_ of 1100% in response to 60 μM Fsk, which was 9−47 times larger than those of other cAMP probes (Fig. 2c-d, Supplementary Fig. 1b). G-Flamp1 also showed large fluorescence increases with a maximum ΔF/F_0_ of 340% and 820% in HeLa and CHO cells, respectively (Supplementary Fig. 12). To rule out possible unspecific responses, we generated a cAMP-insensitive indicator G-Flamp1-mut by introducing the R307E mutation into mlCNBD of G-Flamp1 (Supplementary Fig. 13)^20^. As expected, G-Flamp1-mut showed no detectable signal change in living cells (Fig. 2c). To demonstrate the sensitivity of G-Flamp1, 2.5 nM Iso was exploited to produce a small amount of cAMP in HEK293T cells. G-Flamp1 exhibited an obvious fluorescence increase with a ΔF/F_0_ > 100% after 5 min stimulation while other sensors showed little signal changes (|ΔF/F_0_| < 10%) in our setup (Fig. 2e). Under two-photon excitation (920 nm), G-Flamp1 exhibited a maximum ΔF/F_0_ of 1240%, which is much larger than those of Flamindo2 and cAMPr (−79% and 72%, respectively) (Supplementary Fig. 10b-c). Meanwhile, G-Flamp1 had a 13- and 90-fold higher signal-to-noise ratio (SNR) compared with Flamindo2 and cAMPr, respectively (Supplementary Fig. 10d).

Then we explored the specificity and reversibility of G-Flamp1 in HEK293T cells. Cyclic guanosine monophosphate (cGMP), which is synthesized from guanosine triphosphate (GTP) by guanylyl cyclase in mammalian cells, has been shown to bind cAMP-sensing domains with weaker affinity^14, 31^. To examine the response of G-Flamp1 to cGMP, the sodium nitroprusside (SNP), a nitric oxide (NO) donor that activates soluble guanylyl cyclase, was utilized to induce a large amount of cGMP in living cells. When HEK293T cells were treated with 25 µM SNP, the low-affinity (K_d_ ∼1.09 µM) cGMP sensor Green cGull^32^ showed a maximum ΔF/F_0_ of 210% while G-Flamp1 showed no detectable signal change (Fig. 2f and Supplementary Fig. 14), indicating the high specificity of G-Flamp1 towards cAMP over cGMP. Regarding reversibility, HEK293T cells expressing G-Flamp1 exhibited increased fluorescence upon 100 nM Iso treatment and then returned to basal level after addition of 15 µM β-AR anti-agonist propranolol (Prop) (Fig. 2g).

Besides cell lines, primary cortical neurons were also utilized to examine cellular localization and fluorescence change of G-Flamp1. Again, G-Flamp1 was evenly distributed in neuronal soma and neurites. Upon application of 100 µM AR agonist norepinephrine (NE) or 1 µM Iso, a ΔF/F_0_ of ∼100%−150% was observed in both soma and neurites (Fig. 2h-i). Upon 60 µM Fsk treatment, G-Flamp1 showed significant fluorescence increase with a ΔF/F_0_ of 500%−700% in both soma and neurites (Supplementary Fig. 15). Taken together, G-Flamp1 shows low cytotoxicity, great distribution, decent brightness, large dynamic range and high sensitivity in cell lines and primary neurons at 37 °C.

### *In vivo* two-photon imaging of cAMP dynamics in zebrafish

To test whether G-Flamp1 can function in intact living organisms, we first utilized optically transparent zebrafish embryos under Fsk stimulation. We injected UAS:G-Flamp1(or G-Flamp1-mut)-T2A-NLS-mCherry (nuclear localized mCherry) plasmid into the embryos of EF1α:Gal4 transgenic zebrafish at one-cell stage (Supplementary Fig. 16a). The expression of G-Flamp1 or G-Flamp1-mut sensor was confirmed by green fluorescence in cells of the developing central nervous system. Brain ventricular injection of 120 µM Fsk but not PBS elicited a robust fluorescence increase with a ΔF/F_0_ of 450% for G-Flamp1, whereas no signal changes were observed for G-Flamp1-mut (Supplementary Fig. 16b-d). These data indicate that G-Flamp1 sensor has high sensitivity for *in vivo* cAMP detection in zebrafish.

### *In vivo* two-photon imaging of cAMP dynamics in *Drosophila*

The importance of cAMP in associative learning, where it serves as a coincidence detector by integrating concurrent signal inputs from both conditioned and unconditioned stimuli, has been well documented across phyla^33, 34^. In *Drosophila*, cAMP signaling in the mushroom body (MB) Kenyon cells (KCs) is indispensable for acquiring aversive memory, such as associating specific odor with punitive electrical shock^35, 36^. To reveal cAMP dynamics in living organisms, we generated transgenic flies expressing G-Flamp1 in MB KCs and performed functional two-photon imaging in MB medial lobe (Fig. 3a-c). When the fly was exposed to either 1 s odor puff or subsequent 0.5 s electrical shock, we observed time-locked fluorescence responses with a ΔF/F_0_ of ∼100% (Fig. 3d-e). Compared with the MB β’ lobe that has similar responses among different compartments, the MB γ lobe exhibited compartmentally heterogeneous responses to specific stimuli, as the largest responses were observed in γ4 to odor and in γ2 to electrical shock. These compartmentalized signals were not due to the unequal expression level or saturation of the sensor, since 100 µM Fsk perfusion elicited a homogeneous ΔF/F_0_ of around 250% (Fig. 3f). G-Flamp1 specifically reported cAMP changes since the GFP alone expressed in KCs showed no significant response to 1 s odor, 0.5 s shock or 100 µM Fsk perfusion (Fig. 3d-f). Moreover, both the rise and decay time (τ_on_ and τ_off_) for cAMP changes evoked by odor or shock were similar in different compartments (Fig. 3g-h). Collectively, these results show that G-Flamp1 allows detection of physiologically relevant cAMP dynamics in *Drosophila* with high fidelity and good spatiotemporal resolution, and sheds lights on the role of compartmentally separated cAMP signaling in the olfactory learning process.

**Fig. 3.**
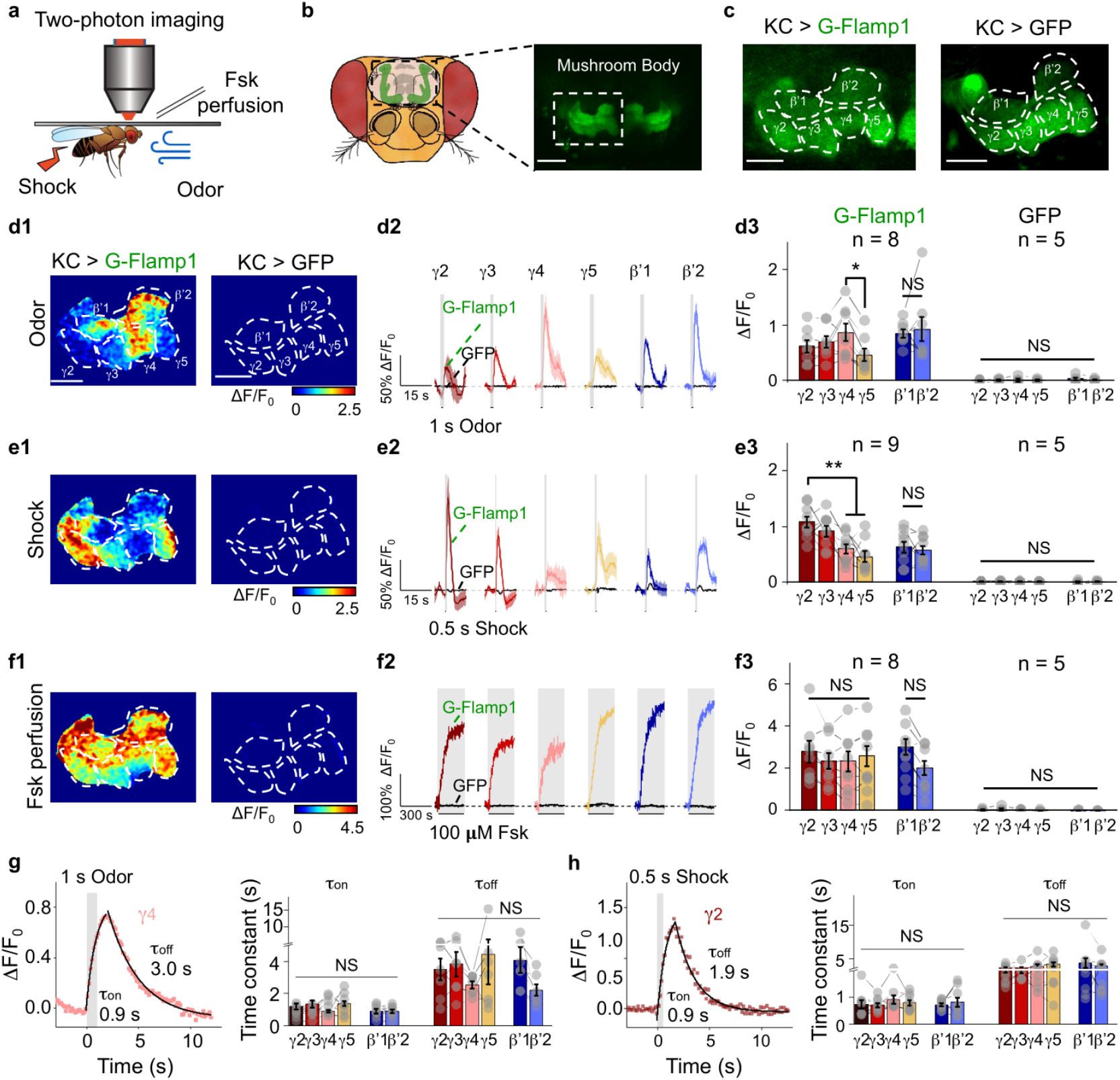
G-Flamp1 reports compartmental cAMP dynamics evoked by physiological stimuli in *Drosophila* through *in vivo* two-photon imaging. (a) Schematics of *in vivo* two-photon imaging setup in *Drosophila* with multiple stimuli. (b) Schematics and fluorescent images of *Drosophila* MB KCs. Scale bar: 50 µm. (c) Fluorescence images of *Drosophila* MB KCs expressing G-Flamp1 (left) or GFP (right). Scale bars: 25 µm. (d) Representative pseudo-color image (d1), traces (d2) and quantification (d3) of fluorescence responses of G-Flamp1 (color) and GFP (black) across MB compartments to 1 s odor. Representative traces were 3-trial average from one fly. n = 8 and 5 for G-Flamp1 and GFP groups, respectively. Two-tailed Student’s *t*-tests were performed in d3. For comparisons of cAMP signals between different MB compartments, *P* = 0.21, 0.37 and 0.048 between γ4 and γ2, γ3 and γ5, respectively. Scale bars: 25 µm. (e) Similar to **d** except that 0.5 s electrical shock was applied to the fly. n = 9 and 5 for G-Flamp1 and GFP groups, respectively. Two-tailed Student’s *t*-tests were performed in e3. For comparisons of cAMP signals between different MB compartments, *P* = 0.368, 0.007 and 0.001 between γ2 and γ3, γ4 and γ5, respectively. (f) Similar to **d** except that 100 µM Fsk was perfused to the fly brain. n = 8 and 5 for G-Flamp1 and GFP groups, respectively. Two-tailed Student’s *t*-tests were performed in f3. *P* > 0.05 between γ2, γ3, γ4 and γ5. (g) Representative traces of ΔF/F_0_ of G-Flamp1 in γ4 evoked by 1 s odor. Data were fitted with single-exponential functions and τ_on_ and τ_off_ values were extracted (left). Quantifications of τ_on_ and τ_off_ for different MB compartments were shown (right). One-way ANOVA test was performed. NS, not significant. (h) Similar to **g** except that 0.5 s electrical shock was applied to the fly. One-way ANOVA test was performed. NS, not significant. Data in **d2**, **e2** and **f2** are shown as mean ± SEM with shaded regions indicating the SEM. Quantifications in **d3**, **e3**, **f3**, **g** (right) and **h** (right) are shown as mean ± SEM overlaid with data points from individual flies. \*\**P* < 0.01, \**P* < 0.05 and NS, not significant.

### *In vivo* two-photon imaging of cAMP dynamics in mouse cortex

To demonstrate the utility of G-Flamp1 sensor to detect physiologically relevant cAMP dynamics in living animals, we performed head-fixed two-photon imaging in the motor cortex (M1) of awake mice during forced locomotion (Fig. 4a), which was reported to be associated with increased neuromodulator and PKA activities^37^. We co-expressed G-Flamp1 (or G-Flamp1-mut) and the red calcium sensor jRGECO1a in the neurons of motor cortex and imaged the layer 2/3 region (Fig. 4b). We observed running-induced, cell-specific, cAMP and calcium signals with no correlation (Fig. 4c). Interestingly, neurons in M1 area could be further divided into three groups based on the cAMP dynamics: ∼60% neurons with fast increase of cAMP (higher average response during the first 30 s after the onset of forced running) and no significant change of calcium, ∼30% neurons with slow increase of cAMP and little change of calcium, and ∼6% neurons with decrease of cAMP and increase of calcium (Fig. 4c). As a control, G-Flamp1-mut showed little fluorescence change (Fig. 4d). Distribution analysis and averaged traces of ΔF/F_0_ of G-Flamp1 and jRGECO1a further confirmed the heterogeneity of neuronal responses (Fig. 4e-i). Therefore, dual-color imaging of calcium and cAMP revealed cell-specific neuronal activity and neuromodulation of cortical neurons in mice during forced locomotion.

**Fig. 4.**
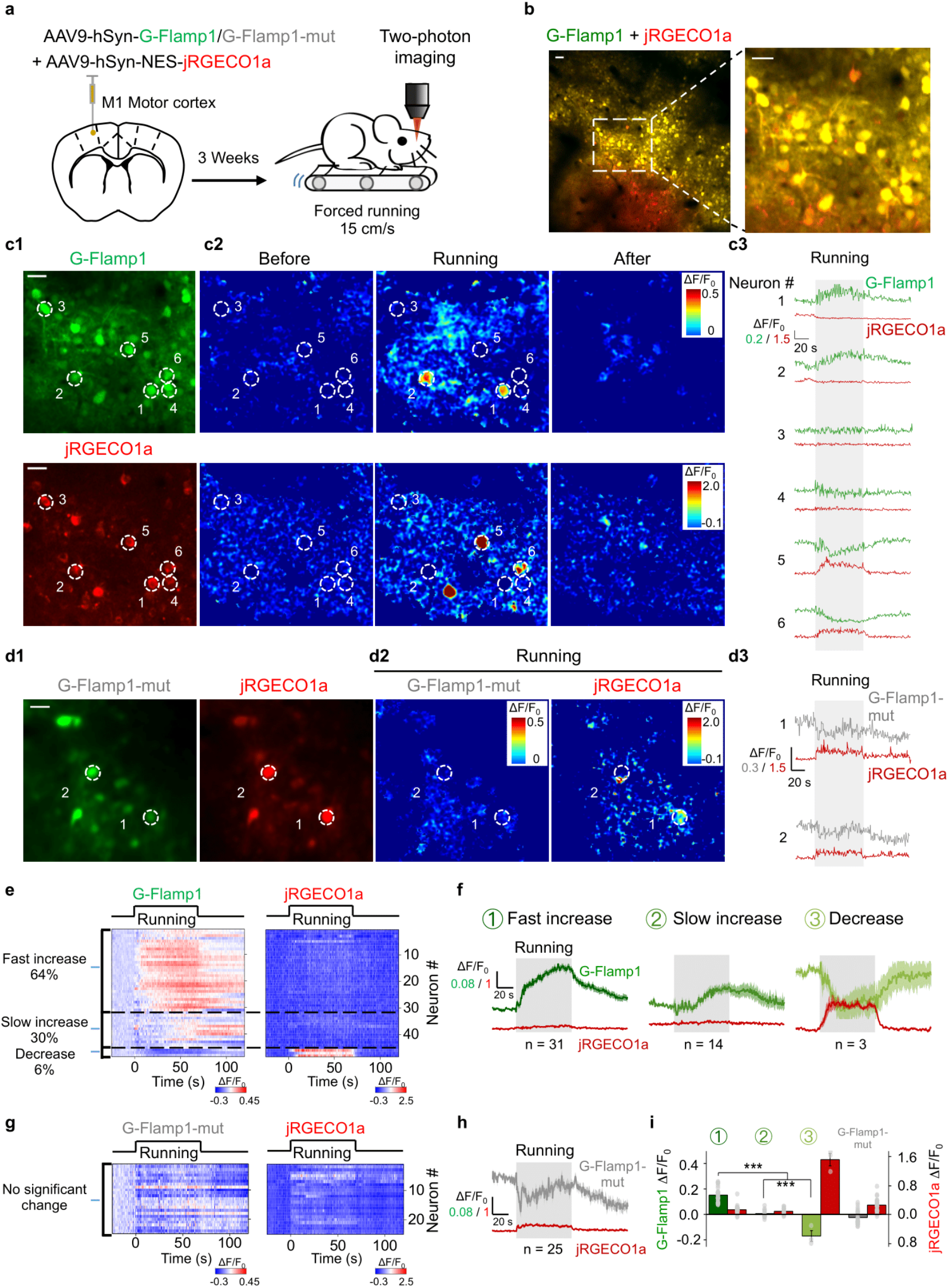
G-Flamp1 reveals forced running-induced cAMP signals of neurons in the mouse motor cortex through *in vivo* two-photon imaging. (**a**) Schematic diagram depicting the head-fixed mice on a treadmill together with two-photon imaging of the motor cortex co-expressing G-Flamp1 (or G-Flamp1-mut) and jRGECO1a. (**b**) Two-photon imaging of the mouse motor cortex co-expressing G-Flamp1 and jRGECO1a. The fluorescence of G-Flamp1 (green) and jRGECO1a (red) was merged and shown in yellow pseudo-color. An ROI (white dashed square with a side length of 260 µm) was selected for following analysis. Scale bars: 50 µm. (**c**) Representative images of G-Flamp1 and jRGECO1a expression in mice (c1), the pseudo-color images (c2) and the traces (c3) of ΔF/F_0_ in response to forced running. White dashed circles with a diameter of 20 µm indicate selected ROIs covering soma for analysis. Scale bars: 30 µm. (**d**) Representative images of G-Flamp1-mut and jRGECO1a expression in mice (d1), the pseudo-color images of ΔF/F_0_ (d2) during the forced running phase and the traces of ΔF/F_0_ (d3) in response to forced running. The white dashed circles with a diameter of 20 µm indicate selected ROIs covering the soma for analysis. Scale bar: 30 µm. (**e**) Heatmaps of G-Flamp1 and jRGECO1a responses during running task. Each row denotes a single cell’s response. n = 48 cells from three mice. (**f**) Averaged traces of ΔF/F_0_ for G-Flamp1 and jRGECO1a for neurons from three groups of different cAMP dynamics. n = 31, 14 and 3 cells for fast increase, slow increase and decrease groups, respectively. **(g)** Heatmaps of G-Flamp1-mut and jRGECO1a responses during running task. Each row denotes a single cell’s response. n = 27 cells from three mice. **(h)** Averaged traces of ΔF/F_0_ for G-Flamp1-mut and jRGECO1a during forced running process. (**i**) Quantification of the average ΔF/F_0_ during the first 30 s after the onset of forced running for G-Flamp1, G-Flamp1-mut and jRGECO1a in **e** and **g**. Two-tailed Student’s *t*-tests were performed. \*\*\**P* < 0.001. Quantifications are shown as mean ± SEM in **f, h** and **i** with shaded regions or error bars indicating the SEM.

### *In vivo* fiber photometry recording of cAMP dynamics in mouse nucleus accumbens

To test the ability of G-Flamp1 sensor to report cAMP dynamics in deep brain regions, we measured cAMP levels in the nucleus accumbens (NAc) using fiber photometry in mice performing a classical conditioning task. The NAc was chosen because it is recently reported that PKA, a downstream molecule in the cAMP signaling pathway, plays a critical role in dopamine-guided reinforcement learning behavior^38^. We first injected an adeno-associated virus (AAV) expressing G-Flamp1 into the NAc and measured fluorescence signals using fiber photometry while the mice were trained to perform the conditioning task (Fig. 5a). In the task, the mice were trained to learn the associations between three auditory cues (conditioned stimulus, CS) and respective outcomes (unconditioned stimulus, US) (Fig. 5b; 8 kHz pure tone → water; white noise → brief air puff to the animal’s face; 2 kHz pure tone → nothing). Well-trained mice had a high licking rate selectively to the water-predictive sound, and the G-Flamp1 signal showed a large increase immediately after the onset of the water-predictive sound, while responses to the other two sounds were much smaller (Fig. 5c-e).

**Fig. 5.**
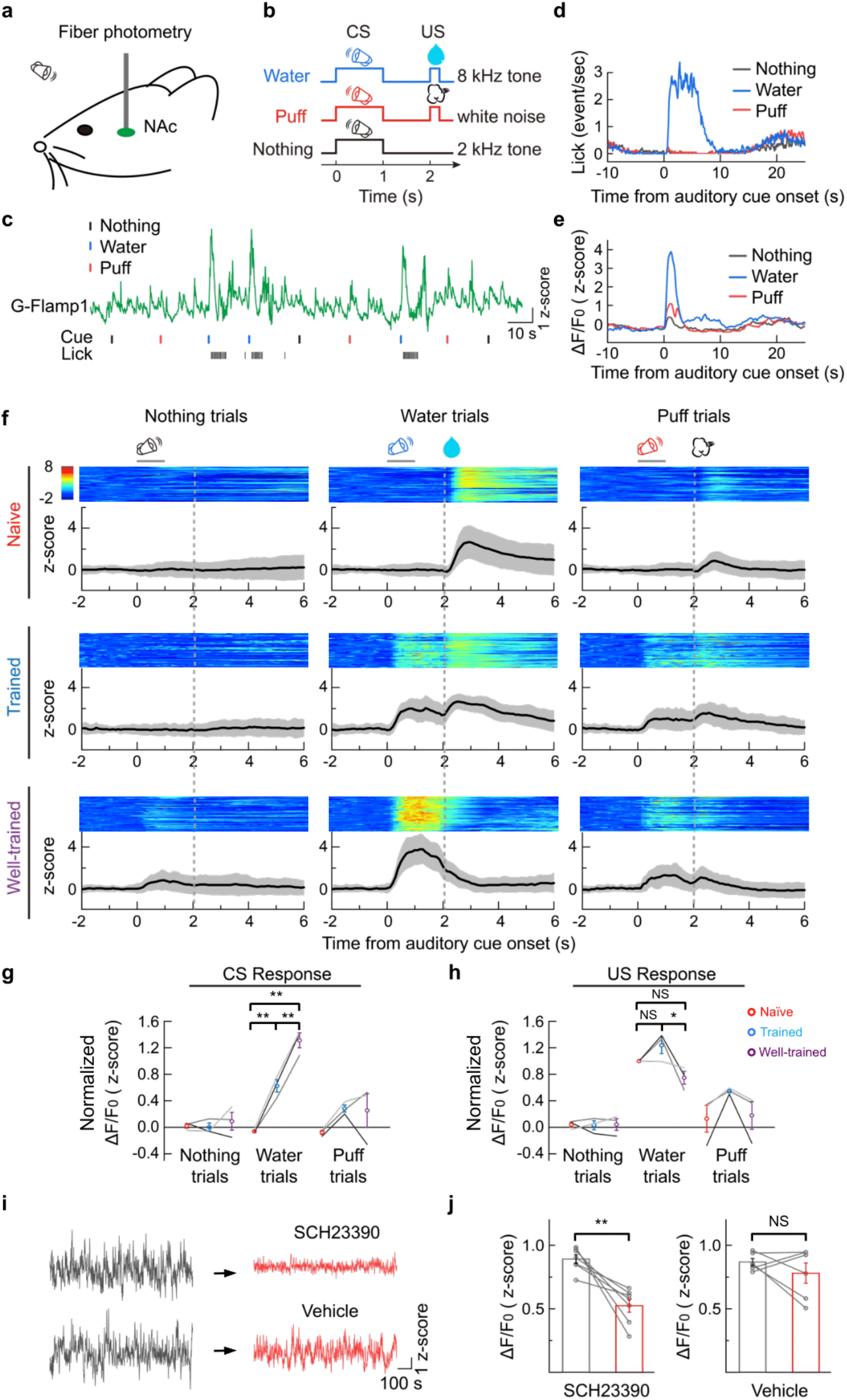
G-Flamp1 reports cAMP activities during an auditory Pavlovian conditioning task in the mouse NAc through *in vivo* fiber photometry. (a) Schematic for fiber photometry recording of G-Flamp1-expressing neurons from the NAc of a head-fixed mouse during an auditory Pavlovian conditioning task. **(b)** Schematic diagrams for the behavioral tasks. The mouse was trained to learn associations between three different auditory cues (conditioned stimulus, CS) and corresponding outcomes (unconditioned stimulus, US). **(c)** Exemplar trace of G-Flamp1 signal from a well-trained mouse encompassing nine sequential trials. The timings of cues (CS) and the lick responses (US) are indicated below. **(d)** Exemplar time-aligned lick responses in **c**. **(e)** Exemplar time-aligned G-Flamp1 signals in **c**. **(f)** Exemplar time-aligned pseudo-color images and averaged traces (mean shaded with ± standard deviation) from a mouse in naïve, trained and well-trained sessions. **(g-h)** Group analysis of the normalized peak Z scores of cAMP signals to CS and US in different sessions. Each trace (coded with specific grey value) represents data from one animal (n = 3 mice). Values with error bars indicate mean ± SEM. Post hoc Tukey’s tests were performed. Water trial CS responses: *P* = 0.00312 between naive and trained, *P* = 6.92772 × 10^-5^ between naive and well-trained, *P* = 0.00312 between trained and well-trained. Water trial US responses: *P* = 0.23198 between naïve and trained, *P* = 0.19808 between naive and well-trained, *P* = 0.02021 between trained and well-trained. **(i)** Exemplar recording of G-Flamp1 signals in NAc before and after injection (i.p.) of D1R antagonist SCH23390 or vehicle. **(j)** Quantification of G-Flamp1 signals before and after SCH23390 (n = 7 recordings from 3 mice, *P* = 0.0038) or vehicle (n = 6 recordings from 3 mice, *P* = 0.34) injection. Two-tailed Student’s *t*-tests were performed in **j**. \*\**P* < 0.01, \**P* < 0.05 and NS, not significant.

Interestingly, the G-Flamp1 signal in the water trials exhibits characteristic dynamics during the learning process: in naïve mice, there was a notable signal increase to water delivery; throughout the training, the magnitude of the water-evoked response decreased, while a response to the reward-predictive sound gradually increased (Fig. 5f-h). This dynamic change mimics the dopamine signal during classical conditioning^39, 40^, suggesting that the increase in cAMP in the NAc is mainly driven by dopamine release. To confirm this, we thus blocked the dopamine D1 receptor using SCH22390 (i.p.) and observed a significantly reduced cAMP signal (Fig. 5i-j). Together, these results demonstrate that the G-Flamp1 sensor has a high signal-to-noise ratio and high temporal resolution to report the dynamic changes of cAMP in behaving mice.

## Discussion

In this study, we described G-Flamp1, a high-performance GEAI engineered by inserting cpGFP into the bacterial mlCNBD. G-Flamp1 exhibits a maximum ΔF/F_0_ of ∼1100% in living cells under both one-photon and two-photon excitation, thus being the most responsive GEAI. We also demonstrated the utility of G-Flamp1 in reporting cAMP dynamics in various model organisms with fiber photometry and optical imaging methods. Given the high sensitivity and direct readout, G-Flamp1 would be useful for screening drugs targeting cAMP signaling pathways using high-content screening assays.

Our *in vivo* two-photon imaging experiments in mouse cortex showed that G-Flamp1 is able to detect bidirectional cAMP changes with single-neuron resolution (Fig. 4). Given that multiple neuromodulators can be released in the motor cortex^37^, different downstream signaling processes are expected to be induced in cortex neurons, which might partially explain the discrepancy between cAMP signal and calcium activity in our results (Fig. 4f). Further studies are needed to dissect out the underlying regulation mechanisms and potential functions. Nevertheless, together with other spectrally compatible sensors, G-Flamp1 will be a useful tool for investigating signal transduction networks in behaving animals.

Very recently, three genetically encoded cAMP indicators (single FP-based Pink Flamindo and cADDis, FRET-based cAMPFIRE-L) have been used for two-photon imaging of cAMP in behaving mice^41–43^, building up the relationship between cAMP signaling and animal behavior. Given its high performance, G-Flamp1 would be an alternative and better choice for *in vivo* cAMP imaging. Compared to GCaMPs, the potential capabilities of G-Flamp1 are only beginning to be realized and will be fully explored in the future. Combined with miniaturized microscopes^44^, G-Flamp1 would be able to visualize cAMP activity patterns in freely moving animals. Moreover, by utilizing G-Flamp1 along with biological models, some long-standing biological questions may be addressed. For example, it may be possible to understand how cAMP is regulated in drug addiction and stress-induced behaviors^45, 46^.

Engineering and structural analysis of G-Flamp1 reveals three interesting findings. First, modest conformation changes of insertion sites in sensing domain can induce large fluorescence change of cpFP. Generally, insertion sites with large structural change are chosen to make large fluorescence change sensors^47^. However, the insertion site in G-Flamp1 is near the mouth of the cAMP-binding pocket and undergoes a small conformational change upon cAMP binding^20^. Second, linkers connecting sensing domain and cpFP can adopt a more rigid conformation. Although random coil and short α-helical turns are observed in single-FP sensors with crystal structures available^26–28^, the first linker along with its flanking sequences in cAMP-bound G-Flamp1 folds as a long β-strand, which may transform into a random coil in the absence of cAMP according to our MD simulation. Third, a different self-contained fluorescence modulation way exists in G-Flamp1. Indeed, similar to G-Flamp1, the fluorescence modulation of red calcium sensors (K-GECO1, R-GECO1 and RCaMP) are also self-contained^26^. However, their chromophores are shielded from bulk solvent in different ways: two linkers along with sensing domain wrap the surface hole of FP caused by circular permutation in red calcium sensors, while only one linker chokes the hole in G-Flamp1. This makes the cpGFP in G-Flamp1 a useful scaffold to be combined with other sensing domains for engineering of new single-FP sensors.

Despite its high performance, G-Flamp1 could be further improved for specific applications. It would be feasible to generate G-Flamp1 variants with improved properties through structure-guided mutagenesis. For example, G-Flamp1 variants with higher basal fluorescence may be useful to monitor cAMP activities in fine structures with high signal-to-background ratio. In addition, G-Flamp1 variants with higher affinity would enable more sensitive detection of subtle changes of cAMP at submicromolar concentration. Besides green G-Flamp1 and its variants, red/near-infrared and photoconvertible sensors using mlCNBD as a sensing domain could be developed to visualize cAMP changes in deep tissue and permanently mark cells with cAMP activities, respectively, which has been realized in calcium sensors^48–50^.

## Methods

### Chemicals and Reagents

cAMP-Na (Cat. No. A6885) and cGMP-Na (Cat. No. G6129) were purchased from Sigma-Aldrich. cAMP (Cat. No. C107047), noradrenaline bitartrate monohydrate (N107258), isoproterenol hydrochloride (Cat. No. I129810) and propranolol (Cat. No. S133437) were purchased from Aladdin (Shanghai, China). Forskolin (Cat. No. S1612) and Enhanced Cell Counting Kit-8 (CCK-8) (Cat. No. C0041) were purchased from Beyotime Biotechnology (Shanghai, China). The CREB antibody 48H2 (Cat. No. 9197S) and phospho-CREB (Ser133) antibody 87G3 (Cat. No. 9198S) were purchased from Cell Signaling Technology, Inc.

### Plasmid construction

Plasmids were generated using the Infusion method (Takara Bio USA, Inc.). PCR fragments were amplified using PrimerStar (normal PCR or site-directed mutagenesis) or Taq (random mutagenesis) DNA polymerases. When needed, overlap PCR was exploited to generate the intact DNA fragment for Infusion. All PCR primers were purchased from Sangon Biotechnology Co., Ltd (Shanghai, China). Plasmids p2lox-cAMPr (Cat. No. 99143), pAAV.Syn.GCaMP6f.WPRE.SV40 (Cat. No. 100837), pAAV.CamKII.GCaMP6s.WPRE.SV40 (Cat. No. 107790) and pAAV.Syn.NES-jRGECO1a.WPRE.SV40 (Cat. No. 100854) were purchased from Addgene. The DNA sequences of Flamindo2, Pink Flamindo, mlCNBD and jRCaMP1b were synthesized by Genscript^11, 13, 16, 51^. pcDNA4-R-FlincA was a gift from Dr. Kazuki Horikawa (Tokushima University). To express fluorescent proteins or sensors in bacterial or mammalian cells, cDNAs of FPs or sensors were subcloned into pNCS or pCAG vector^52^, respectively. To improve G-Flamp1’s stability in mammalian cells, its N-terminal arginine immediately after the initiator methionine was deleted^53^. cDNAs of G-Flamp1, G-Flamp1_opt_ and G-Flamp1-mut_opt_ (opt: mouse/human codon optimized) were subcloned into AAV vectors to make AAV2-CAG-G-Flamp1, AAV2-hSyn-G-Flamp1 and AAV2-hSyn-G-Flamp1-mut. pCAG-mEGFP and pCAG-mCherry were kept in our lab. All constructs were confirmed by DNA sequencing (Sangon Biotechnology Co., Ltd, Shanghai, China).

### Screening of cAMP sensors expressed in bacteria

Two mlCNBD fragments (Gly213-Pro285 and Asn286-Ala355) and cpGFP with linkers from GCaMP6f were amplified, overlapped and cloned into BamHI/EcoRI sites of pNCS vector with an N-terminal 6×His tag for protein purification. Site-directed and random mutagenesis were performed via overlap PCR and error-prone PCR, respectively. The DNA libraries were transformed into DH5α cells lacking adenylate cyclase gene *CyaA* (DH5α-*ΔCyaA*), which were generated by the phage λ Red recombination system^54^. After overnight incubation at 34°C, colonies with different fluorescence intensities on the LB agar plates were screened by eye in a BlueView Transilluminator (Vernier) with the 400 nm-500 nm excitation light and a yellow acrylic long-pass filter, or by fluorescence imaging in a home-made imaging system with 480/20 nm excitation and 520/20 nm emission filters. To quantitatively compare the brightness of selected variants, bacterial patches on the agar plates cultured overnight at 34 °C were: 1) imaged in the home-made system mentioned above and analyzed by ImageJ software (National Institutes of Health) (Supplementary Fig. 3d and f), or 2) collected in PBS and the OD_600_-normalized fluorescence intensities were measured with an Infinite M1000 fluorometer (Tecan) (Supplementary Fig. 9).

The fluorescence changes of cAMP sensors in response to cAMP were examined using the bacterial lysate. Briefly, selected bacterial colonies were patched on LB agar plate and grew at 25°C for 3 days. The harvested bacterial cells were suspended in 1 mL of HEPES buffer (150 mM KCl and 50 mM HEPES-KOH, pH 7.15) and lysed by sonication followed by centrifugation. 120 µL of clear lysates were mixed with 2 µL of HEPES buffer or 2 µL of 30 mM cAMP or 2 µL of 30 mM cGMP and then the fluorescence were recorded with an Infinite M1000 PRO fluorometer (Tecan). The fluorescence change ΔF/F_0_ was calculated as (F-F_0_)/F_0_, where F and F_0_ are fluorescence intensities of sensors in the presence or absence of cAMP (or cGMP), respectively.

### Bacterial protein expression, purification and *in vitro* characterization

DH5α-*ΔCyaA* cells were transformed with pNCS-FP or sensor and cultured overnight at 34 °C. The colonies were then patched on LB agar plates and cultured at room temperature for 3 days. The harvested bacterial cells were suspended in HEPES buffer and lysed by sonication. His-tagged recombination proteins were purified with cobalt-chelating affinity chromatography (Pierce) and desalted with HEPES buffer (pH 7.15) using the gel filtration column (Bio-Rad).

Quantum yields were determined using mEGFP as a standard (QY = 0.60). Extinction coefficients were determined according to the ‘base denatured chromophore’ method^52^. pH titrations were performed using a series of pH buffers ranging from 2 to 10.5 (50 mM Citrate-Tris-Glycine buffer. The desired pH was achieved by adding 2 M of sodium hydroxide or 2 M of hydrochloric acid)^52^. The fluorescence excited at 450 nm in different pH buffers was measured using an Infinite M1000 PRO fluorometer. The fluorescence intensities were plotted against the pH values and the pKa was determined by fitting the data to the Henderson-Hasselbalch equation^55^.

To determine the affinity of G-Flamp1, 1 µM of purified protein in HEPES buffer was mixed with varying concentrations of cAMP (0.001, 0.01, 0.1, 0.5, 1, 2, 5, 10, 25, 100 and 500 µM) or cGMP (0.01, 0.1, 0.5, 1, 2, 5, 10, 25, 100, 500, 1000 and 2000 µM). The fluorescence excited at 450 nm were recorded with an Infinite M1000 PRO fluorometer. The fluorescence change ΔF/F_0_ was plotted against the cAMP or cGMP concentrations and fitted by a sigmoidal binding function to determine the K_d_ and Hill coefficient^49^.

The association constant (k_on_) and dissociation constant (k_off_) between G-Flamp1 and cAMP were determined using Chirascan spectrometer equipped with an SX20 Stopped-Flow accessory (Applied Photophysics Ltd). Briefly, 1.6 µM of protein solution was mixed 1:1 with cAMP of different concentrations (0.5, 1, 2, 5, 10 and 50 µM) and the fluorescence excited at 480 nm were measured with a 520/30 nm filter. The data were fitted using the following single-exponential function^56, 57^: F(t) = F_0_ + A_obs_ × exp(-k_obs_ × t), where F(t) is the value of fluorescence increase at time t, F_0_ is the final value of fluorescence increase, A_obs_ is the amplitude of the exponentially decreasing part and k_obs_ is the observed first-order rate constant. The k_on_ and k_off_ were fitted using the following equation: k_obs_ = k_on_ × [cAMP] + k_off_, where [cAMP] is the concentrations of cAMP used. The association and dissociation half-time t_on_ and t_off_ were calculated as ln2/(k_on_ × [cAMP]) and ln2/k_off_, respectively.

To get the excitation wavelength-dependent brightness and ΔF/F_0_ under two-photon excitation, purified proteins were excited with wavelengths from 700 to 1000 nm with a 20 nm step size on a Nikon-TI two-photon microscope equipped with a Ti:sapphire laser and a 25 × 1.4 NA water immersion objective. The 495-532 nm fluorescence were collected and the intensities were then normalized to laser powers at different wavelengths.

### Crystallization and structure determination of G-Flamp1

The coding sequence of G-Flamp1 was cloned into pSUMO expression vector with 6× His and SUMO tags at the N-terminus. *E.coli* BL21 (DE3) pLysS cells were transformed with pSUMO-G-Flamp1 and grew on LB agar overnight at 34°C. Colonies were expanded in LB media at 34°C and induced at OD 0.6 with 0.1 mM IPTG for additional 3 hours at 34°C. The harvested cells were lysed with a high-pressure homogenizer at 1000 bar in binding buffer (20 mM Imidazole, 500 mM NaCl, 20 mM Tris-HCl, pH 7.5). The protein was purified on a Ni Sepharose 6 Fast Flow column (GE Healthcare) under gravity and eluted with the elution buffer (300 mM Imidazole, 500 mM NaCl, 20 mM Tris-HCl, pH 7.5). The elution was incubated with ULP1 protease and dialyzed against the dialysis buffer (100 mM NaCl, 10 mM β-ME, 20 mM Tris-HCl, pH 7.5) overnight at 4°C and purified again on a Ni Sepharose 6 Fast Flow column to remove the 6×His and SUMO tags and ULP1 protease. After concentration, the flow-through was loaded on a Hiload 16/600 Superdex 200 pg column (GE Healthcare) in the dialysis buffer for further purification. Fractions containing purified protein were pooled, concentrated and incubated with cAMP at 1:5 molar ratio for 1 hour at 4°C. Crystals were grown using the hanging drop vapour diffusion method with 2 µL protein solution (10 mg/mL) and 2 µL reservoir solution (40% v/v PEG 400, 100 mM Imidazole, pH 8.0). The mixture was equilibrated against 300 µL reservoir solution at 20°C for 5 days. Crystals were flash-frozen for X-ray diffraction data collection. A data set was collected to 2.2 Å resolution at wavelength 1.0000 Å on beamline BL17B1 of the Shanghai Synchrotron Radiation Facility (SSRF). Data sets were processed with HKL3000^58^. The structure was solved by molecular replacement method using Phaser software^59^ implanted in the Phenix program suite^60^, with cpGFP (PDB: 3EVP) and mlCNBD (PDB: 3CLP) as search models. The model building was performed manually using the Coot^61^.

### Molecular dynamics simulations

The X-ray crystal structure of cAMP-bound G-Flamp1 was first modified by removing cAMP, water molecules and solvent ions. With the AMBER14 force field in YASARA version 19.9.12.L.64^62^, the modified structure was subjected to molecular dynamics (MD) simulations in a box of 85.43 Å × 72.66 Å × 69.53 Å dimensions containing 12734 water molecules. MD simulation conditions were as follows: 1 bar of pressure, 298 K of temperature, pH 7.4 and 1 fs time step. The MD run was a 157.50 ns length with snapshots taken every 100 ps. The trajectory was analyzed by YASARA and the simulated model was considered in the equilibrium state. The last snapshot was converted to a PDB file for further analysis. A movie was also produced by YASARA for visualizing the continuous conformation change of Trp75 during the MD run.

### Cell culture, DNA transfection and virus infection

Mammalian cell lines were maintained in DMEM (HEK293T and HeLa cells) or DMEM/F12 (CHO cells) supplemented with FBS (10% v/v) and penicillin/streptomycin (both at 100 units/mL) in a humidified incubator at 37°C with 5% CO_2_. Plasmid transfections of cultured cells were performed according to the Lipofectamine 2000 protocol. Primary cortical neurons were prepared from embryonic day 16 (E16) BALB/c mice as previously described^63^ and kept in Neurobasal medium with B27 (2%) and penicillin/streptomycin (both at 100 units/mL). DIV (days *in vitro*) 7-9 neurons were infected with AAV8-CAG-G-Flamp1 virus prepared using PEG8000/NaCl solution and imaged at DIV13-18.

### Stable cell line generation and proliferation rate measurement

The CAG promoter and G-Flamp1 was inserted between two terminal inverted repeats for *piggyBac* transposase (PBase) in pPB-LR5 vector^64^ to make pPB-LR5-CAG-G-Flamp1. HEK293T cells in a 24-well plate were co-transfected with 1 µg of pCMV-hyperactive PBase^64^ and 1 µg of pPB-LR5-CAG-G-Flamp1, expanded for 1 week and then sorted for medium-brightness ones with a BD FACSAria III Cell Sorter (BD, USA). The proliferation rates of HEK293T control cells or cells expressing G-Flamp1 were measured using the Enhanced Cell Counting Kit-8 (Cat. No. C0041, Beyotime Biotechnology, Shanghai, China).

### Western blotting

Total protein of cells were extracted by radioimmunoprecipitation assay (RIPA) buffer (Beyotime Biotechnology, Shanghai, China) and protein concentrations were measured using BCA Protein Assay kit (Pierce, USA). Equal amounts of protein were separated by 4%-10% SDS-PAGE, transferred on PVDF membranes, and immuno-detected with primary antibodies against pCREB and CREB. Signal detection was carried out on a ChemiDoc MP imaging system (Bio-Rad) using the ECL kit (Cat. No. #32106, Pierce, USA).

### Wide-field fluorescence imaging of cAMP indicators in living cells

Wide-field imaging was performed on an Olympus IX83 microscope equipped with a 63 × 1.4 numerical aperture (NA) objective (HEK293T, HeLa and CHO cells) or a 20 × 0.75 NA objective (cultured neurons). Briefly, mammalian cells grown on glass-bottom dishes (Cat. No. #FD35-100, World Precision Instruments) were transfected with indicated plasmids and 24 hours later serum-starved for 2-4 hours. The culture medium was replaced with live cell imaging solution right before fluorescence imaging. Time-lapse images were captured every 15 s. The excitation and emission filters used for different sensors were as follows: ex 480/30 nm and em 530/30 nm for green sensors (GCaMP6s, cAMPr, Flamindo2 and G-Flamp1), ex 568/20 nm and em 630/50 nm for red sensors (jRCaMP1b, Pink Flamindo and R-FlincA), ex 441/20 nm and em 530/30 nm for G-Flamp1. Background-subtracted fluorescence was used to calculate fluorescence change ΔF/F_0_ that is defined as (F-F_0_)/F_0_, where F_0_ is the baseline signal before stimulation.

### Two-photon fluorescence imaging of cAMP indicators in living cells

Two-photon imaging was performed on a Nikon-TI two-photon microscope equipped with a Ti:sapphire laser and a 25 × 1.4 NA water immersion objective. In brief, mammalian cells grown on glass-bottom dishes were transfected with indicated plasmids and 24 hours later serum-starved for 2-4 hours. The culture medium was replaced with live cell imaging solution right before fluorescence imaging. Cells were excited with a 920 nm laser line and detected via a 495-532 nm filter. Time-lapse images were taken every 5 s. Background-subtracted fluorescence intensity was used to calculate ΔF/F_0_. The SNR was defined as the ratio of peak ΔF/F_0_ to the standard deviation of the basal fluorescence before stimulation.

### Brightness comparison of cAMP indicators in HEK293T cells

Fluorescent intensity of indicators was measured using an Infinite M1000 fluorometer or optical microscope. For fluorometer, HEK293T cells grown in 12-well plates were transfected with pCAG-G-Flamp1, pCAG-cMAPr, pCAG-Flamindo2, pCAG-Pink Flamindo, pCAG-R-FlincA, pCAG-GCaMP6s, pCAG-jRCaMP1b, pCAG-mEGFP or pCAG-mCherry construct separately using Lipofectamine 2000. 48 hours later, the cells were washed once with PBS, suspended in live cell imaging solution (Cat. No. A14291DJ, Invitrogen) and transferred to a clear flat-bottom 96-well plate. The fluorescence was recorded under 480 nm excitation. For wide-field or two-photon microscopy, HEK293T cells on glass-bottom dishes were transfected with indicated constructs using Lipofectamine 2000. 48 hours later, the culture medium was replaced with live cell imaging solution and fluorescence images were taken under 480/30 nm (one-photon) or 920 nm (two-photon) excitation.

### Two-photon imaging in zebrafish

cDNAs of G-Flamp1 (or G-Flamp1-mut) and NLS-mCherry (nuclear localized mCherry) were subcloned into pTol2-UAS vector to make pTol2-UAS:G-Flamp1 (or G-Flamp1-mut)-T2A-NLS-mCherry, where T2A is a self-cleaving peptide. Plasmids above with Tol2 mRNA were co-injected into EF1α:Gal4 embryos at one-cell stage. At 52 hours post-fertilization, the brain ventricle of larval zebrafish was injected with PBS or 120 µM Fsk and imaged with a BX61WI two-photon microscope (Olympus) equipped with a 25 × 1.05 NA water immersion objective. The excitation wavelength was 960 nm and 495-540 nm fluorescence was collected. The fluorescence intensities of cells pre- and post-treatment were extracted using ImageJ. Fluorescence change was calculated as ΔF/F_0_, where F_0_ was the average intensity before treatment.

### Two-photon imaging of transgenic flies

The coding sequence of G-Flamp1 was cloned into pJFRC28 (Addgene plasmid #36431). The vector was injected into embryos and integrated into attP40 via phiC31 by the Core Facility of Drosophila Resource and Technology (Shanghai Institute of Biochemistry and Cell Biology, Chinese Academy of Sciences). Stock 30Y-Gal4 (III) is a gift from Yi Rao lab (Peking University). Stock UAS-GFP (III) is a gift from Donggen Luo lab (Peking University). Flies UAS-G-Flamp1/+; 30Y-Gal4/+ and UAS-GFP/30Y-Gal4 were used. Flies were raised on standard cornmeal-yeast medium at 25°C, with 70% relative humidity and a 12 h/12 h light/dark cycle.

Adult females within 2 weeks post-eclosion were used for *in vivo* imaging with a two-photon microscope FV1000 (Olympus) equipped with the Mai Tai Ti:Sapphire laser (Spectra-Physics) and a 25 × 1.05 NA water immersion objective (Olympus). The excitation wavelength was 930 nm and a 495−540 nm emission filter was used. The sample preparation was similar as previously described^40^. Before and after odor stimulation, 1000 mL/min constant pure air was applied to the fly. During 1 s odor stimulation, 200 mL/min air containing isoamyl acetate (Cat. No. 306967, Sigma-Aldrich) mixed with 800 mL/min pure air was delivered to the fly. For electrical shock, 80 V 500 ms electrical stimulus was applied to the fly via copper wires attached to the abdomen. For Fsk application, the blood-brain barrier was carefully removed and Fsk was applied with a 100 µM final concentration. Customized Arduino code was used to synchronize the imaging and stimulation protocols. The sampling rate during odor stimulation, electrical shock stimulation and Fsk perfusion was 6.7 Hz, 6.7 Hz and 1 Hz, respectively.

### Animals

All procedures for animal surgery and experimentation were conducted using protocols approved by the Institutional Animal Care and Use Committees at Shenzhen Institute of Advanced Technology-CAS, Peking University and Institute of Neuroscience-CAS.

### Two-photon imaging in mice

AAV9-hSyn-G-Flamp1, AAV9-hSyn-G-Flamp1-mut and AAV9-hSyn-NES-jRGECO1a viruses were packaged at Vigene Biosciences (Jinan, China). Wild-type female C57 BL/6J mice (6-8 weeks old) were anesthetized with an injection of Avertin or isoflurane (3% induction; 1–1.5% maintenance). The skin and skull above the motor cortex were retracted from the head and a metal recording chamber was affixed. ∼300 nL of AAV was injected into the motor cortex (AP, 1.0 mm relative to bregma; ML, 1.5 mm relative to bregma; depth, 0.5 mm from the dura). A 2 mm × 2 mm or 4 mm × 4 mm square coverslip was used to replace the skull. Three weeks after virus injection, wake mice were habituated for about 15 min in the treadmill-adapted imaging apparatus to minimize the potential stress effects of head restraining. The motor cortex at a depth of 100-200 µm below the pial surface was imaged using a Bruker Ultima Investigator two-photon microscope equipped with the Spectra-Physics Insight X3 and a 16 × 0.8 NA water immersion objective. 920 nm laser line was used for excitation of both green and red indicators. 490-560 nm and 570-620 nm filters were used for green and red fluorescence collection, respectively. The sampling rate was 1.5 Hz. For imaging analysis, we first corrected motion artifact using motion correction algorism (EZcalcium)^65^ and bleed-through between green and red channels using the spectral unmixing algorithm (see details in https://imagej.nih.gov/ij/plugins/docs/SpectralUnmixing.pdf). The fluorescence intensities of ROIs covering the somata were extracted using ImageJ software. Background-subtracted fluorescence intensity was used to calculate ΔF/F_0_.

### Fiber photometry recording of cAMP signals in behaving mice

The AAV9-hSyn-G-Flamp1 virus was packaged at Vigene Biosciences (Jinan, China). Virus was unilaterally injected into NAc of adult C57BL/6N mice (male, > 8 weeks old). During the surgery, mice were deeply anesthetized with isoflurane (RWD Life Science) and mounted on a stereotaxic apparatus (RWD Life Science). Approximately 300 nL of AAV2/9-hSyn-G-Flamp1 (titer 7.29 × 10^13^, 1:7 diluted with 1× PBS before use) was injected into the NAc (AP, + 1.0 mm; ML, + 1.5 mm; −3.9 mm from cortical surface) at a speed of 23 nL/injection (inter-injection interval 15-30 s) using a microinjection pipette injector (Nanoject II, Drummond Scientific). A 200 µm optic fiber (Thorlabs, FT200UMT) housed in a ceramic ferrule was implanted to the same coordinate two weeks later and a stainless steel headplate was affixed to the skull using machine screws and dental cement. After recovery (> 5 days), the mouse was water-restricted to achieve 85-90% of normal body weight and prepared for behavior training. Mice were trained on an auditory conditioning task, in which three auditory cue -outcome pairs (or CS-US pairs; 8 kHz pure tone → 9 µL water; white noise → brief air puff on face; 2 kHz pure tone → nothing) were randomly delivered with 10-20 second randomized inter-trial intervals. The duration of each sound is 1 second and sound intensity was calibrated to 70 dB. The outcomes were delivered 1 second after offset of each sound. The behavioral setup consisted of a custom-built apparatus allowing head fixation of mice. Licking behavior was detected when the tongue of the mouse contacted the water delivery tube. Lick signal was processed in an Arduino UNO board with custom code and sent digitally to the training program (written in Matlab) via a serial port. Water delivery was precisely controlled by a stepping motor pump and air puff (15 psi, 25 ms) was controlled by a solenoid valve. Timing of the pump and valve was controlled by the same Arduino UNO board used for lick detection, which also provides synchronization between the training program and data accusation system (RZ2, TDT). During first two days of each training, the outcomes were delivered without the prediction cues. To record the fluorescence signal from the cAMP sensor, an optic fiber (Thorlabs, FT200UMT) was attached to the implanted ferrule via a ceramic sleeve. The photometry rig was constructed using parts from Doric Lens, which includes a fluorescence optical mini cube (FMC4_AE(405)_E(460-490)_F(500-550)_S), a blue led (CLED_465), a led driver (LED_2) and a photo receiver (NPM_2151_FOA_FC). During recording, a software lock-in detection algorithm (modulation frequency: 459 Hz; low-pass filter for demodulated signal: 20Hz, 6th order) was implemented in a real-time processor (RZ2 with fiber photometry gizmo in Synapse software). The intensity of excitation light was measured as ∼70 µW from tip of the optical fiber. The photometry data was stored using a sampling frequency of 1017 Hz. To analyze the recording data, we first binned the raw data to 10.17 Hz (down-sampled by 100), fitted the binned data with a 2^nd^ order exponential function using Matlab Curve Fitting Tool. The fitting data was then subtracted from the binned data in order to remove the baseline drift resulting from photo-bleaching, and baseline corrected data was converted to z-score for further analysis. To analyze CS-or US-evoked changes in cAMP signals, we aligned each trial to the auditory cue onset and calculated the peri-stimulus time histogram (PSTH). To compare PSTH changes during different phases of the training, we used data from the 2^nd^ day as naïve, the 5^th^ day as trained and 11^th^ day as well-trained. Response to CS was defined as peak of the PSTH between CS onset to US onset and response to US was calculated accordingly using data from US onset to 2 seconds after US onset. To examine the contribution of dopamine signaling to the cAMP signals in NAc during spontaneous wakefulness, a potent dopamine receptor antagonist, SCH23390 (ab120597, Abcam; 0.2 mg/kg in 100 µL 0.9 % NaCl, i.p.) was administered to mice after tens of minutes of baseline was recorded. To be noted, recordings were not interrupted during the i.p. injection. Each mouse used for analysis had been administered with both SCH23390 and vehicle (100 µL 0.9% NaCl, i.p.), but only one of the solutions was used each single day. To quantify the change in cAMP signals, we take the mean of the z-score transformed signal to get Fig. 5j.

### Statistics

The statistical significances between groups were determined using two-tailed Student’s *t*-tests, One-way ANOVA tests (Fig. 3g-h) or Post hoc Tukey’s tests (Fig. 5g-h) with OriginPro 9.1 (OriginLab). \**P* < 0.05, \*\**P* < 0.01, \*\*\**P* < 0.001 and NS (not significant) for *P* > 0.05.

## Data availability

The atomic coordinates and structure factors of the G-Flamp1 (no RSET peptide) and cAMP complex have been deposited in the Protein Data Bank (http://www.rcsb.org) with PDB ID code 6M63. All G-Flamp1 plasmids will be deposited into Addgene (http://addgene.org). The DNA coding sequence of G-Flamp1 will be deposited in Genbank (https://www.ncbi.nlm.nih.gov/genbank). The protein sequence of G-Flamp1 has been provided in Supplementary Fig. 4c. All source data will be provided upon request.

## Code availability

The custom Arduino code for stimulation and two-photon imaging in *Drosophila*, the custom MATLAB and Arduino codes for fiber photometry in mice, and the custom MATLAB code for data analysis will be provided upon request.

## Acknowledgements

This work was supported by National Key Research and Development Program of China (2020YFA0908802, 2017YFA0700403), National Natural Science Foundation of China (81927803, 21874145, 32000732, 32000731), Guangdong Basic and Applied Basic Research Foundation (2020B121201010), Natural Science Foundation of Shenzhen (JCYJ20200109115633343), CAS grants (DWKF20200001, NSY889021058). We thank Drs. Michael Lin at Stanford University, Jonathan A. Cooper at Fred Hutchinson Cancer Research Center, François St-Pierre at Baylor College of Medicine, Yu Mu at Institute of Neuroscience-CAS, Yulin Zhao at Peking University and Fang Liu at the Campbell Family Mental Health Research Institute, for the critical reading of this manuscript.

## Author Contributions

J.C. conceived and supervised the study. L.W. and J.C. designed the study. L.W. performed experiments related to the development and characterization of G-Flamp1 *in vitro*, in cell lines and isolated neurons with help from W.L., Y.C., Y.G., Y.L. and P.T. C.L. performed western blot experiments. Z.Z. performed protein crystallization experiments. Y.Z. and M.H. performed molecular dynamics simulations. S.Y. performed two-photon imaging experiments in zebrafish. J.Z., Y.Y. and X.L. performed two-photon imaging experiments in flies. C.W. performed two-photon imaging experiments in behaving mice. W.P. performed fiber photometry recording experiments in behaving mice. All authors contributed to data interpretation and analysis. L.W., M.X., Y.L. and J.C. wrote the manuscript with input from other authors.

## Competing Interests

The authors declare no competing interests.

## Supplementary Information

### 1. Supplementary Figures

**Supplementary Fig. 1.**
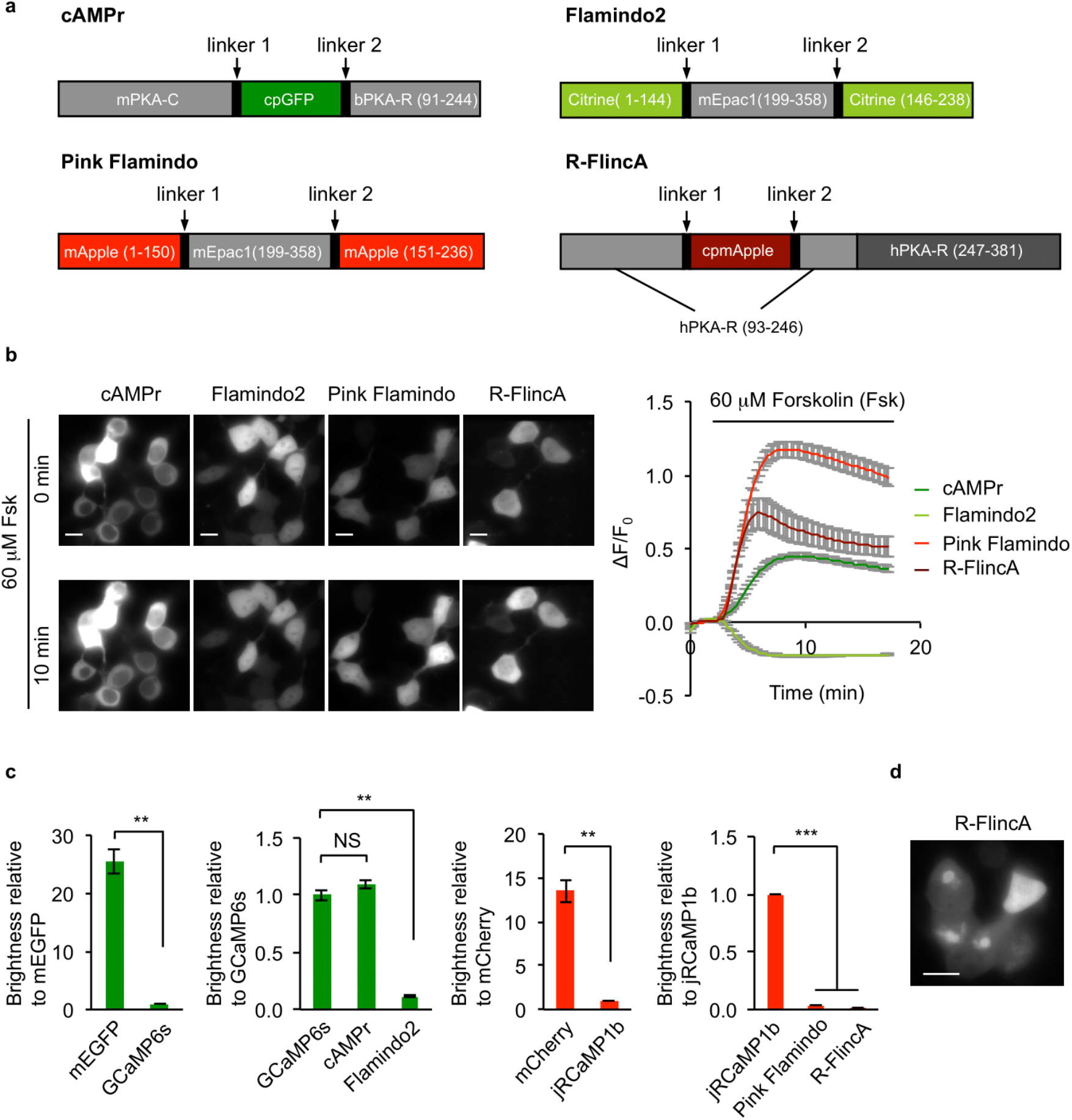
Key properties of previously reported single FP-based cAMP sensors in HEK293T cells at 37°C. (a) Schematic of cAMPr, Flamindo2, Pink Flamindo and R-FlincA sensors. PKA-C and PKA-R represent PKA catalytic and regulatory subunit, respectively. mPKA, bPKA and hPKA are mouse, bovine and human PKA, respectively. mEpac1 is mouse Epac1. GFP, Citrine and mApple are fluorescent proteins. In R-FlincA, cpmApple was inserted into the first CNBD of PKA-R. (b) Representative fluorescence images (left) and traces of ⊗F/F_0_ (right) of cAMP sensors in response to 60 µM Forskolin (Fsk) in HEK293T cells. Notably, the image contrasts for different sensors were different to render fluorescence visible. Data are shown as mean ± SEM. n = 33 cells (cAMPr), 42 cells (Flamindo2), 34 cells (Pink Flamindo) and 18 cells (R-FlincA) from 3 cultures for each sensor. Scale bars: 10 µm. (c) Baseline brightness of the green cAMP sensors (cAMPr and Flamindo2) and red cAMP sensors (Pink Flamindo and R-FlincA) in HEK293T cells. Brightness of green and red cAMP sensors were normalized to those of the green calcium sensor GCaMP6s and the red calcium sensor jRCaMP1b, respectively. The brightness of GFP and mCherry were also normalized to GCaMP6s and jRCaMP1b, respectively. Data are shown as mean ± SEM. n = 3 wells from 12-well plates for each sensor. Two-tailed Student’s *t*-tests were performed. *P* = 0.008 between mEGFP and GCaMP6s, *P* = 0.213 between GCaMP6s and cAMPr, *P* = 0.002 between GCaMP6s and Flamindo2. *P* = 0.0097 between mCherry and jRCaMP1b, *P* = 1.0 × 10^-6^ between jRCaMP1b and Pink Flamindo2, *P* = 5.6 × 10^-7^ between jRCaMP1b and R-FlincA. (d) R-FlincA formed puncta in HEK293T cells after 48 h transfection. Scale bar: 10 µm. \*\*\**P* < 0.001, \*\**P* < 0.01 and NS, not significant.

**Supplementary Fig. 2.**
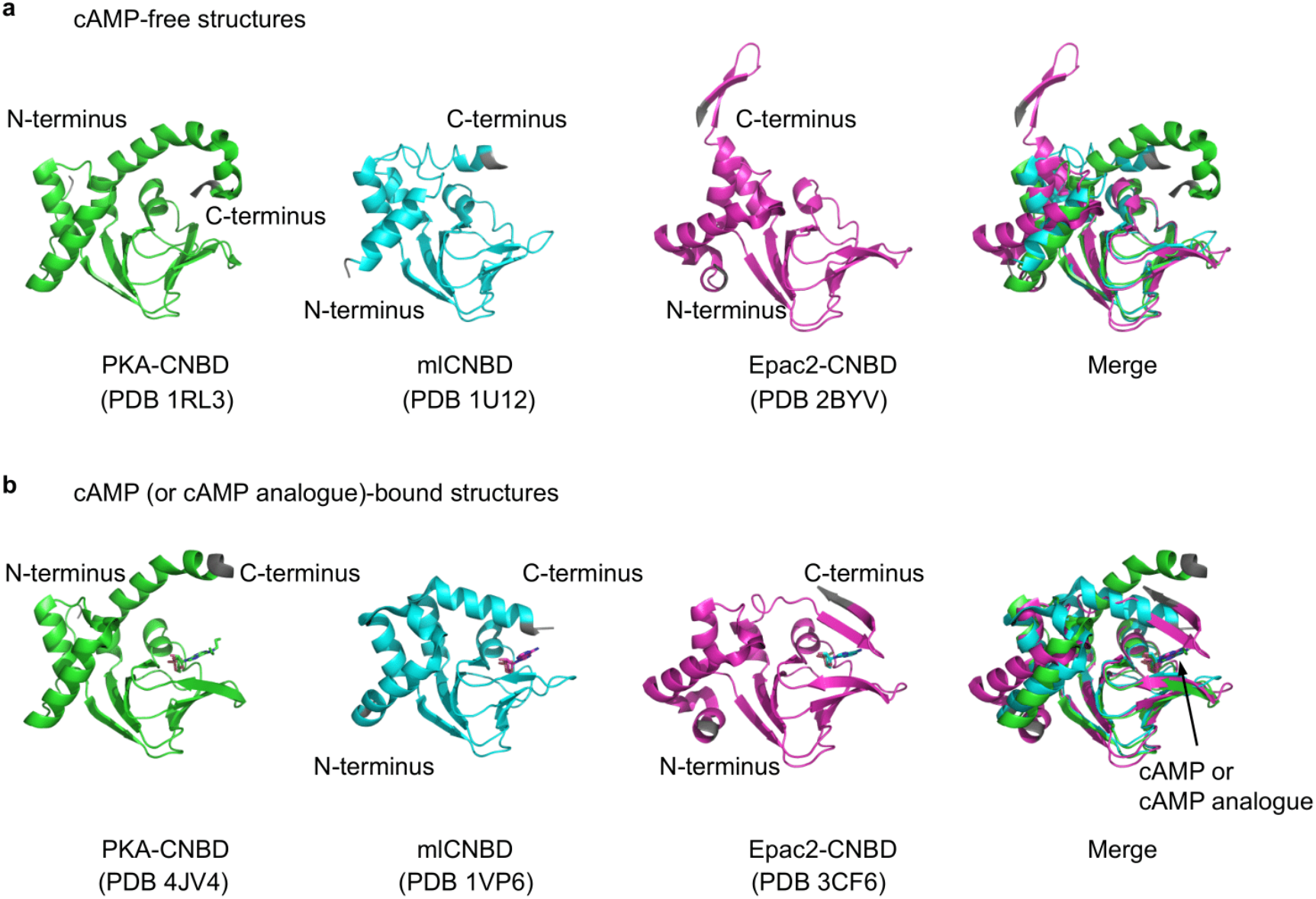
Structure alignments of CNBDs from bovine PKA, mouse Epac2 and bacterial MlotiK1 channel. (a) Structures of different cAMP-free CNBDs. (b) Structures of different cAMP (or its analogue)-bound CNBDs. cAMP or its analogue molecules are shown as stick models. Protein termini are highlighted in grey.

**Supplementary Fig. 3.**
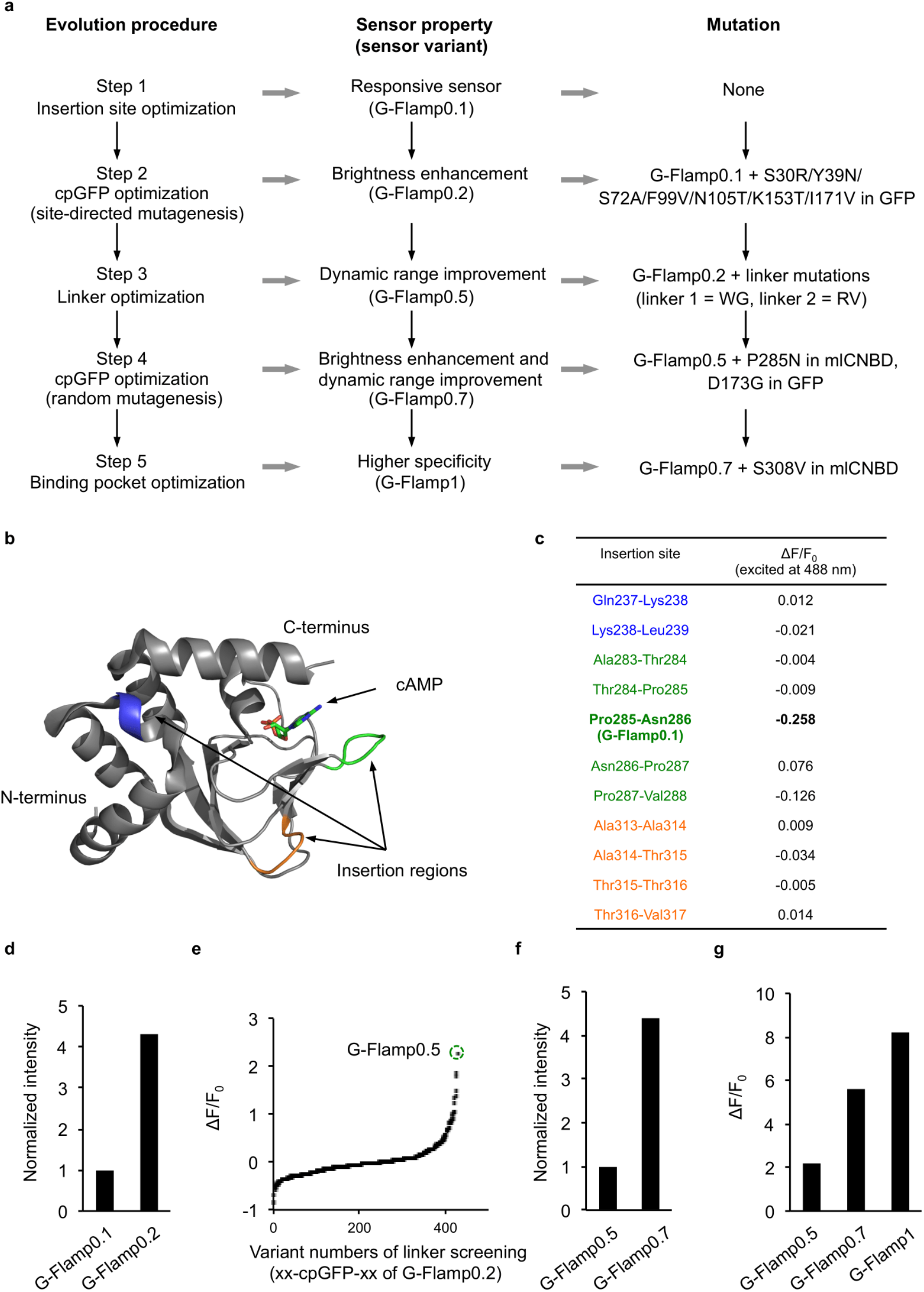
Evolution of G-Flamp1. (a) Five-step directed evolution procedure of G-Flamp1. (b) Three insertion regions tested are highlighted in blue, green and orange in mlCNBD’s structure (PDB 1VP6). (c) ΔF/F_0_ of 11 G-Flamp variants with different insertion sites in response to 500 µM cAMP. The variant with the insertion site between Pro285 and Asn286 (named G-Flamp0.1) showed the largest fluorescence change. The color coding matches the one in **b**. (d) The brightness of G-Flamp0.1 and G-Flamp0.2 in bacterial cells cultured overnight at 34°C. (e) ΔF/F_0_ of 427 G-Flamp0.2 variants with different linkers in response to 500 µM cAMP. The variant with the linkers ‘WG’ and ‘RV’ (named G-Flamp0.5) showed the greatest fluorescence change. (f) The brightness of G-Flamp0.5 and G-Flamp0.7 in bacterial cells cultured overnight at 34°C. (g) ΔF/F_0_ of G-Flamp0.5, G-Flamp0.7 and G-Flamp1 under excitation at 488 nm.

**Supplementary Fig. 4.**
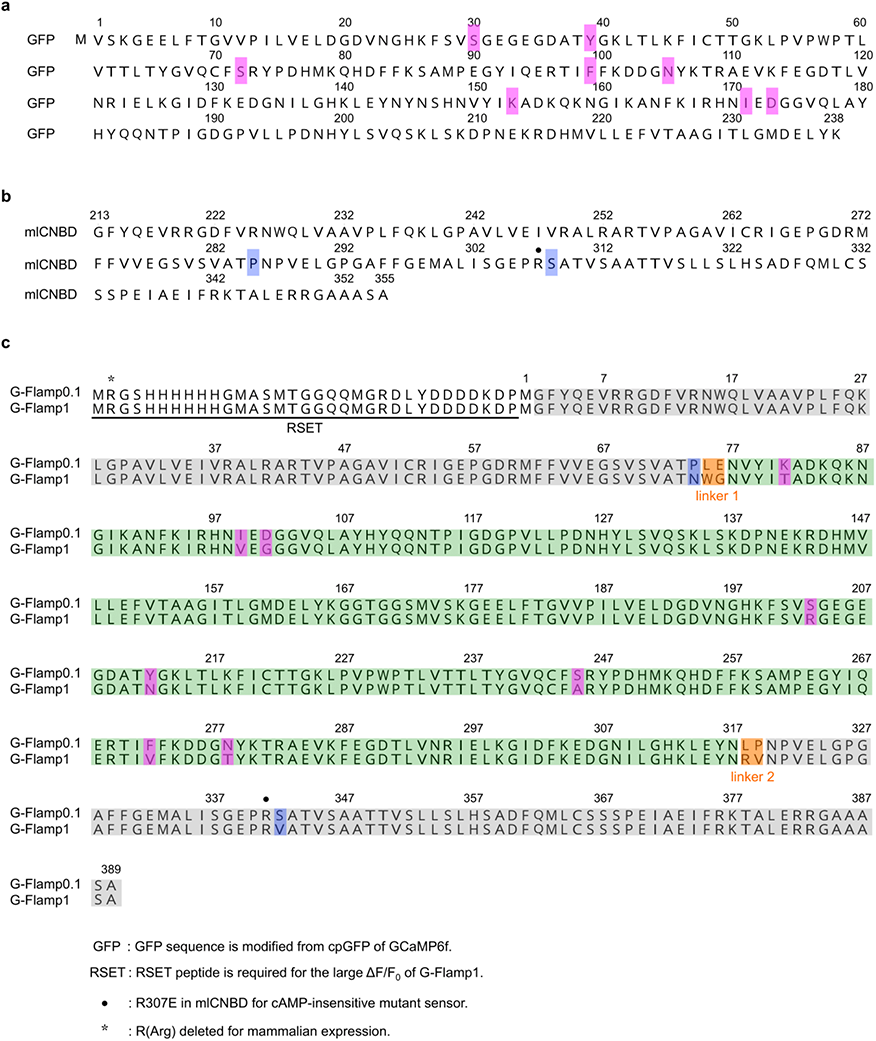
Protein sequences of GFP, mlCNBD, G-Flamp0.1 and G-Flamp1. (a-b) Protein sequences of GFP and mlCNBD. The numberings of GFP and mlCNBD are according to PDB 2Y0G and 1VP6, respectively. Modified amino acid residues in G-Flamp1 sensor are highlighted in magenta and blue. (c) Sequence alignment of full-length G-Flamp0.1 and G-Flamp1. The numbering is according to PDB 6M63. Modified amino acid residues are highlighted in magenta, blue and orange. Note the amino acid Arg immediately after the initiator methionine in G-Flamp1 was deleted for mammalian expression.

**Supplementary Fig. 5.**
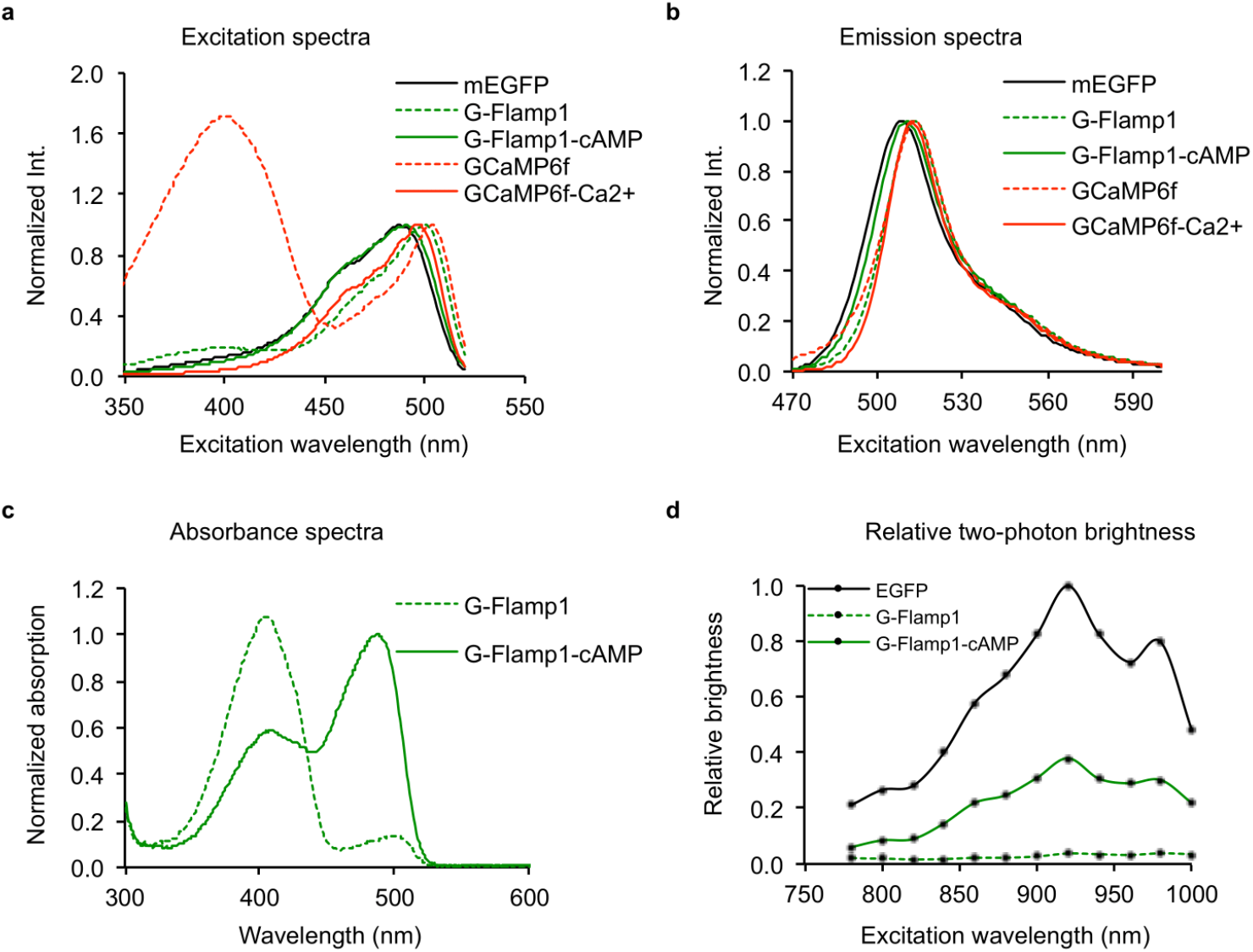
Fluorescence and absorption spectra of purified G-Flamp1, mEGFP and GCaMP6f. (a-b) Excitation (a) and emission (b) spectra of mEGFP, cAMP-free G-Flamp1, cAMP-bound G-Flamp1, calcium-free GCaMP6f and calcium-bound GCaMP6f. (c) Absorption spectra of 20 µM purified G-Flamp1 in HEPES buffer in the presence or absence of 500 µM cAMP. (d) Relative brightness at different excitation wavelengths under two-photon excitation.

**Supplementary Fig. 6.**
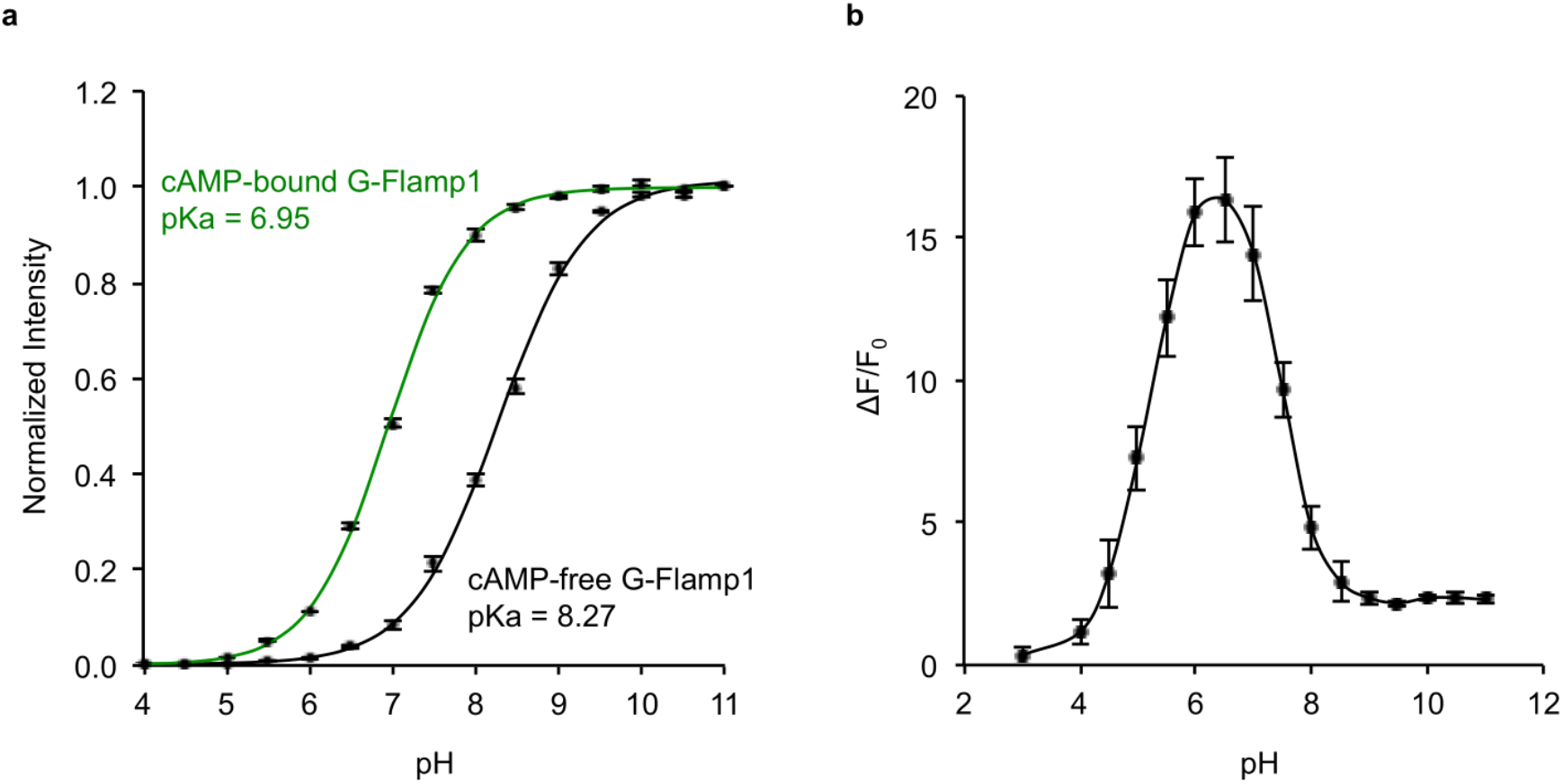
pH-dependent fluorescence and fluorescence change of purified G-Flamp1. (a) Normalized fluorescence of purified G-Flamp1 (2 µM) at various pH values in the presence or absence of 500 µM cAMP. Fitted data are shown as solid lines. (b) ΔF/F_0_ of purified G-Flamp1 (2 µM) in buffers with different pH values. Data are presented as mean ± SEM. n = 3 independent experiments.

**Supplementary Fig. 7.**
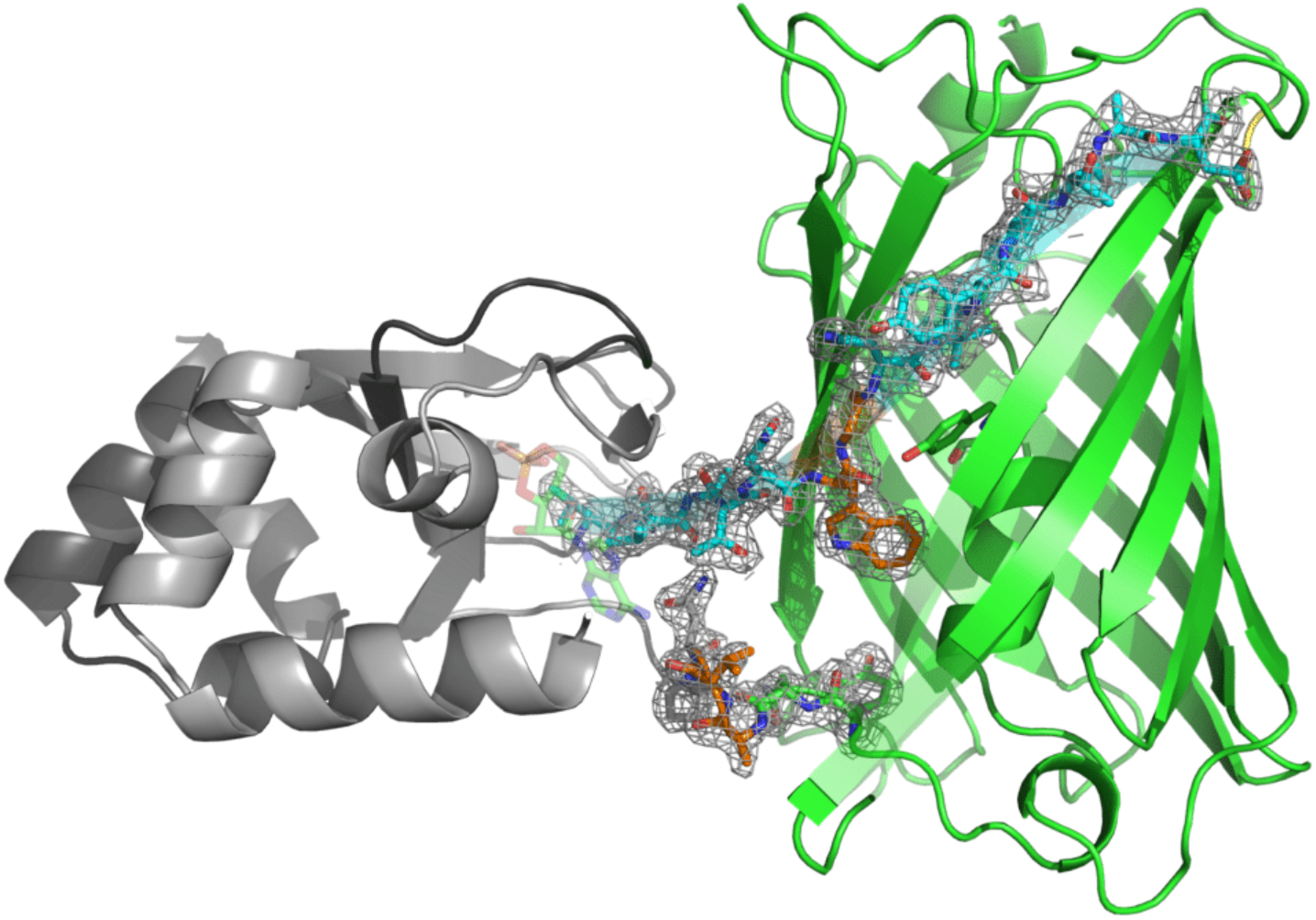
Crystal structure of cAMP-bound G-Flamp1 (PDB: 6M63) with electron density on both two linkers and their neighboring residues. The mesh depicts electron density in the 2Fo−Fc map contoured to 1.2 sigma within 2.0 Å of the atoms displayed in stick form.

**Supplementary Fig. 8.**
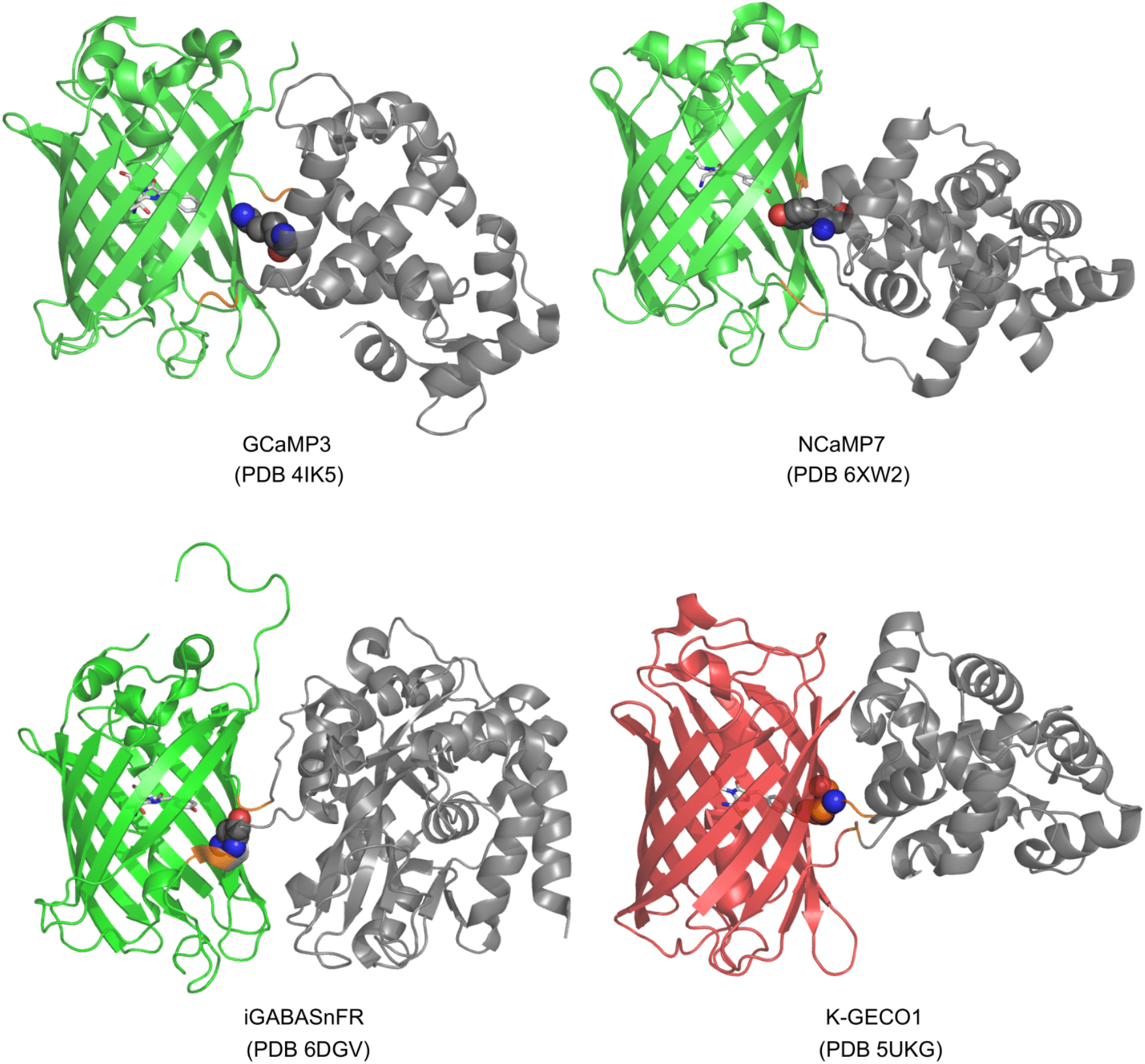
Linker conformation and the interactions between key residues and chromophore in other single-FP indicators. FPs, linkers and sensing domains are marked in green/red, orange and grey, respectively. All chromophores in the FP are shown as stick and amino acid residues interacting with the phenolic oxygen of the chromophore are shown as sphere. In iGABASnFR, the linker 2 folds as α -helix. Unlike GCaMP3 and NCaMP7, in which the fluorescence modulation is dependent on the interactions with residues of CaM, the fluorescence change in K-GECO1 is mediated by a residue from linker 1.

**Supplementary Fig. 9.**
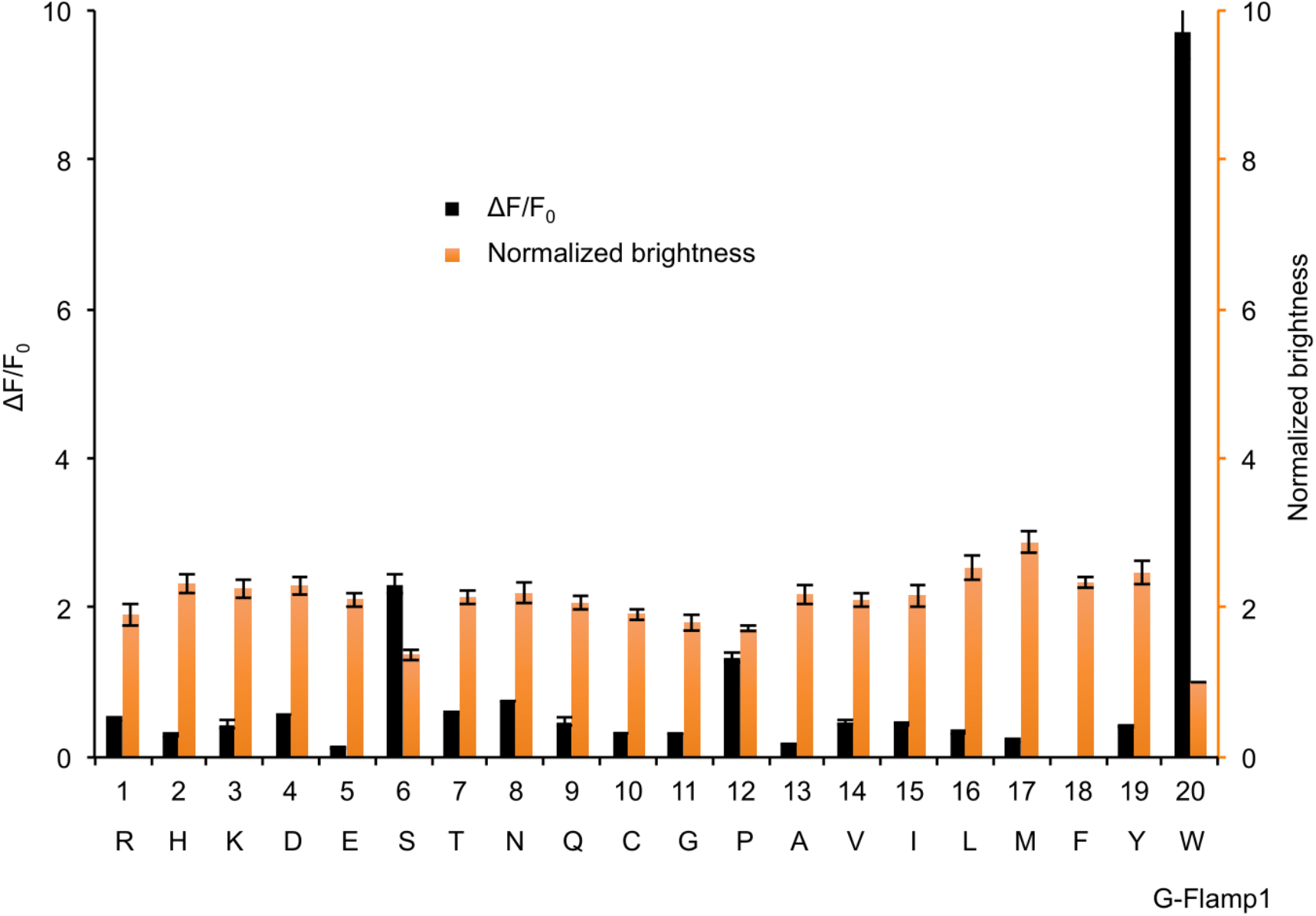
Saturation mutagenesis of Trp75 in G-Flamp1 sensor. Basal brightness (orange) and ΔF/F_0_ (black) for each variant were shown. Error bars indicate SEM of the mean from 3 independent experiments.

**Supplementary Fig. 10.**
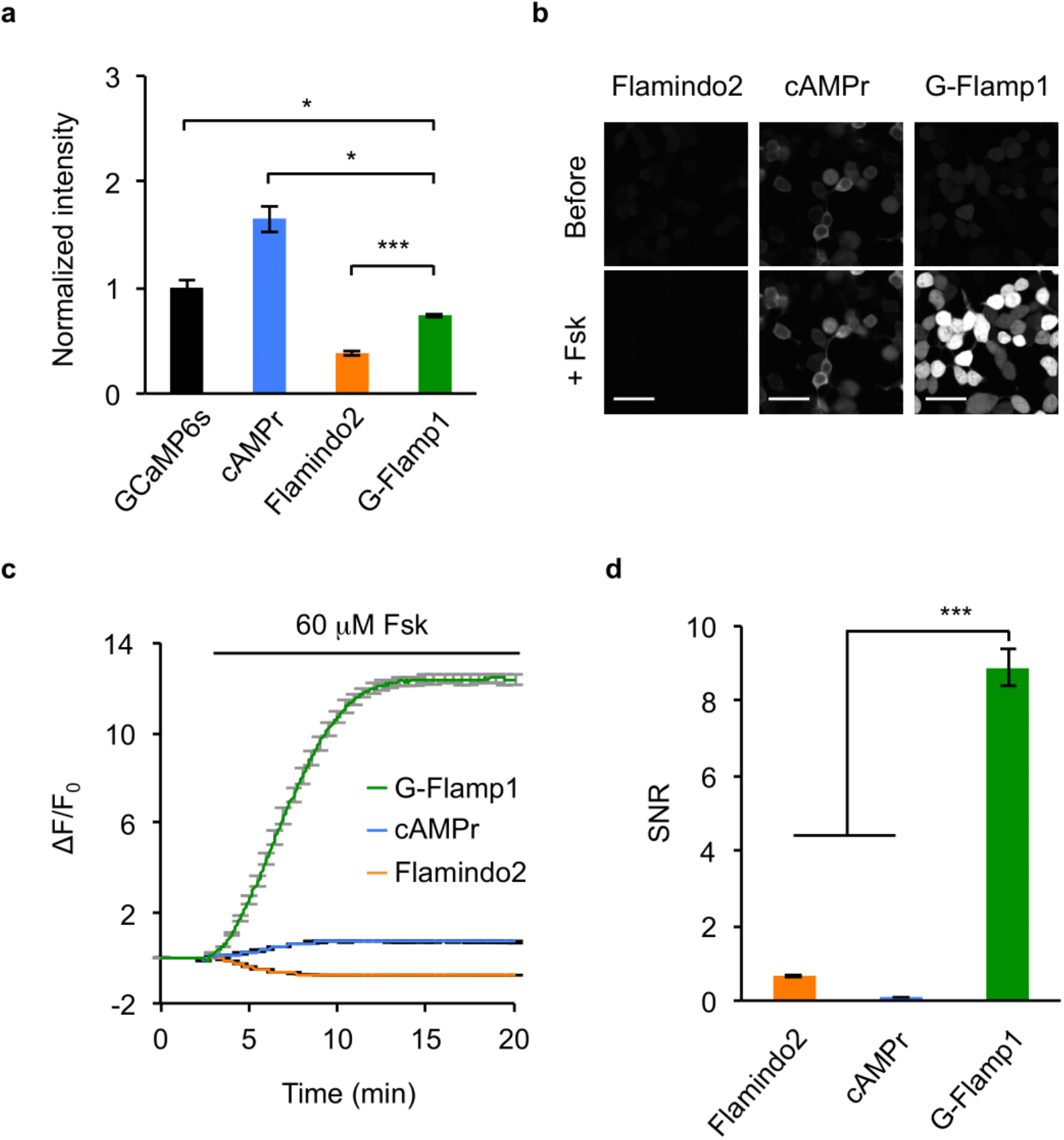
Performance of G-Flamp1 in HEK293T cells under two-photon imaging. (a) Brightness comparison of three different green cAMP sensors (cAMPr, Flamindo2 and G-Flamp1) and GCaMP6s. Images were taken after 48 hours transfection under two-photon excitation (920 nm). n = 3 cultures for each sensor. Two-tailed Student’s *t*-tests were performed. *P* = 0.044, 0.017 and 1.9 × 10^-4^ between G-Flamp1 and GCaMP6s, cAMPr and Flamindo2, respectively. (b-c) Representative two-photon fluorescence images (b) and traces of ΔF/F_0_ (c) of HEK293T cells expressing cAMP sensors in response to 60 µM Fsk. n = 76 cells (Flamindo2), 35 cells (cAMPr) and 64 cells (G-Flamp1) from 2 separate experiments. Scale bars: 50 µm. (d) Signal-to-noise ratio (SNR) of different sensors in (c). Two-tailed Student’s *t*-tests were performed. *P* = 1.1 × 10^-28^ between G-Flamp1 and Flamindo2, and *P* = 1.1 × 10^-26^ between G-Flamp1 and cAMPr. All data are shown as mean ± SEM in **a, c** and **d.** \*\*\**P* < 0.001 and \**P* < 0.05.

**Supplementary Fig. 11.**
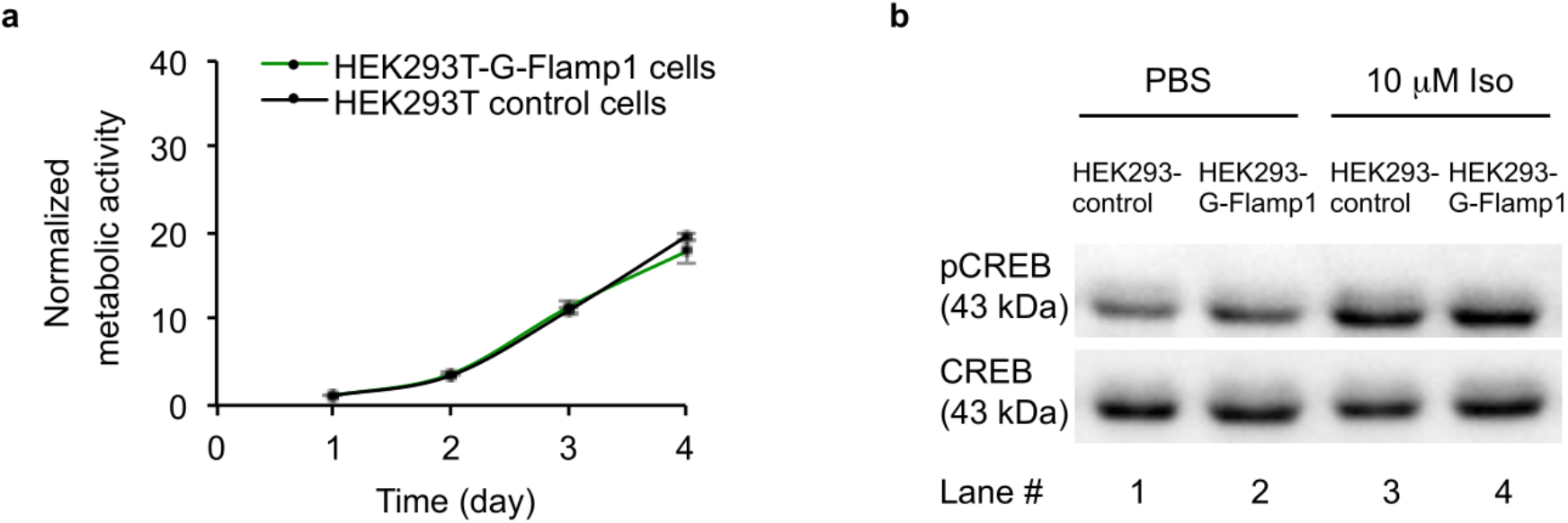
Effects of G-Flamp1 expression on HEK293T proliferation and cAMP signaling. (a) Proliferation rates of HEK293 cells (control) and stable HEK293T cells expressing G-Flamp1 (stable) were measured using the CCK-8 assay. Data are shown as mean ± SEM from 3 independent experiments. (b) Western blot analysis of phosphorylated CREB (pCREB) in cells induced by 10 µM Iso. Representative images from 3 separate experiments are shown. Lanes 1 and 2 were the lysate of serum-starved control and stable HEK293T cells, respectively. Lanes 3 and 4 were the lysate of control and stable HEK293T cells stimulated by 10 µM Iso for 1 hour, respectively.

**Supplementary Fig. 12.**
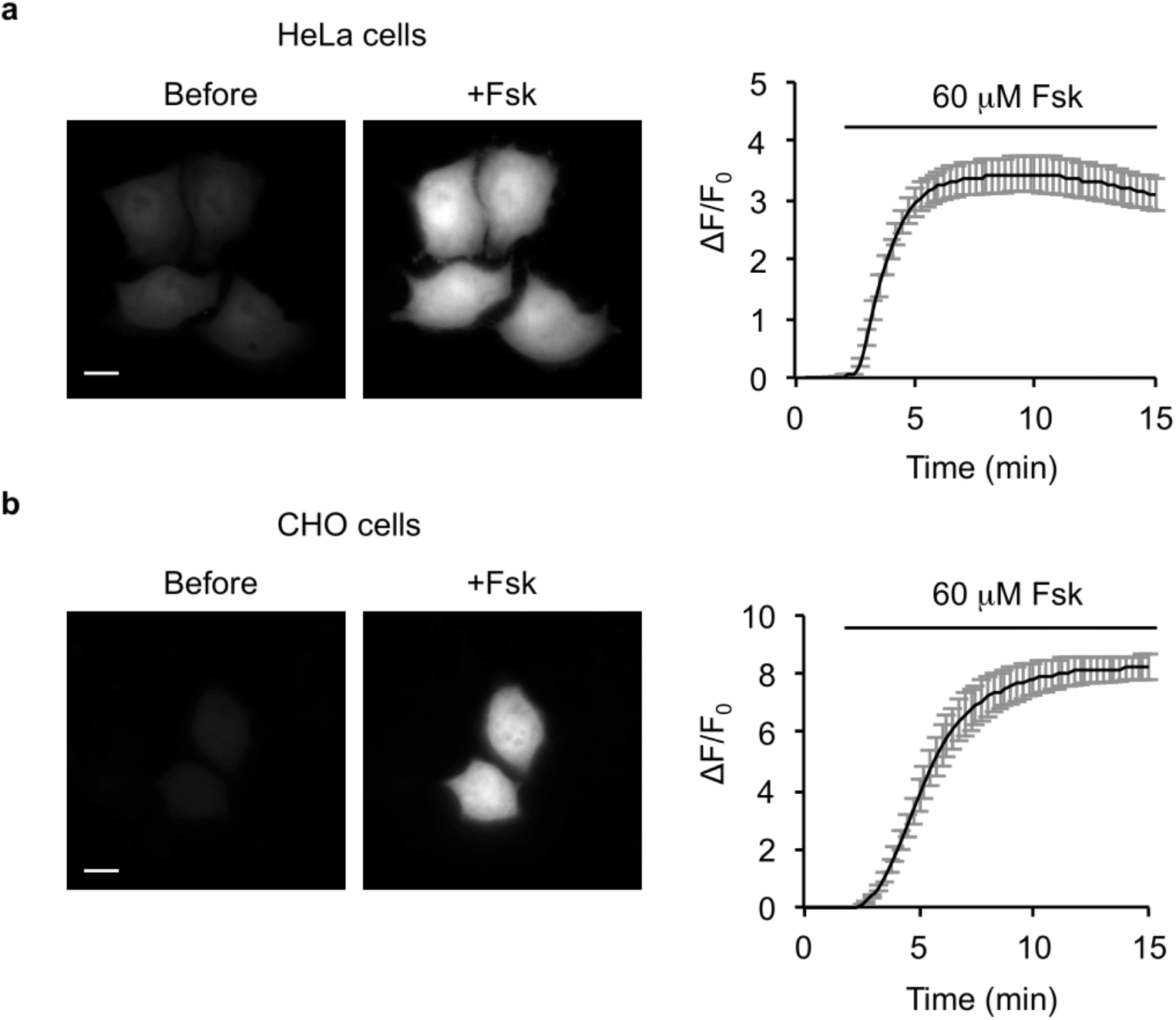
ΔF/F_0_ of G-Flamp1 in HeLa and CHO cells. (a) Representative fluorescence images (left) and ΔF/F_0_ traces (right) of HeLa cells expressing G-Flamp1 in response to 60 µM Fsk. n = 18 cells from 2 cultures. (b) Same as **a** except that the mammalian cell line used was CHO. n = 13 cells from 6 cultures. Data are shown as mean ± SEM. Scale bars: 10 µm.

**Supplementary Fig. 13.**
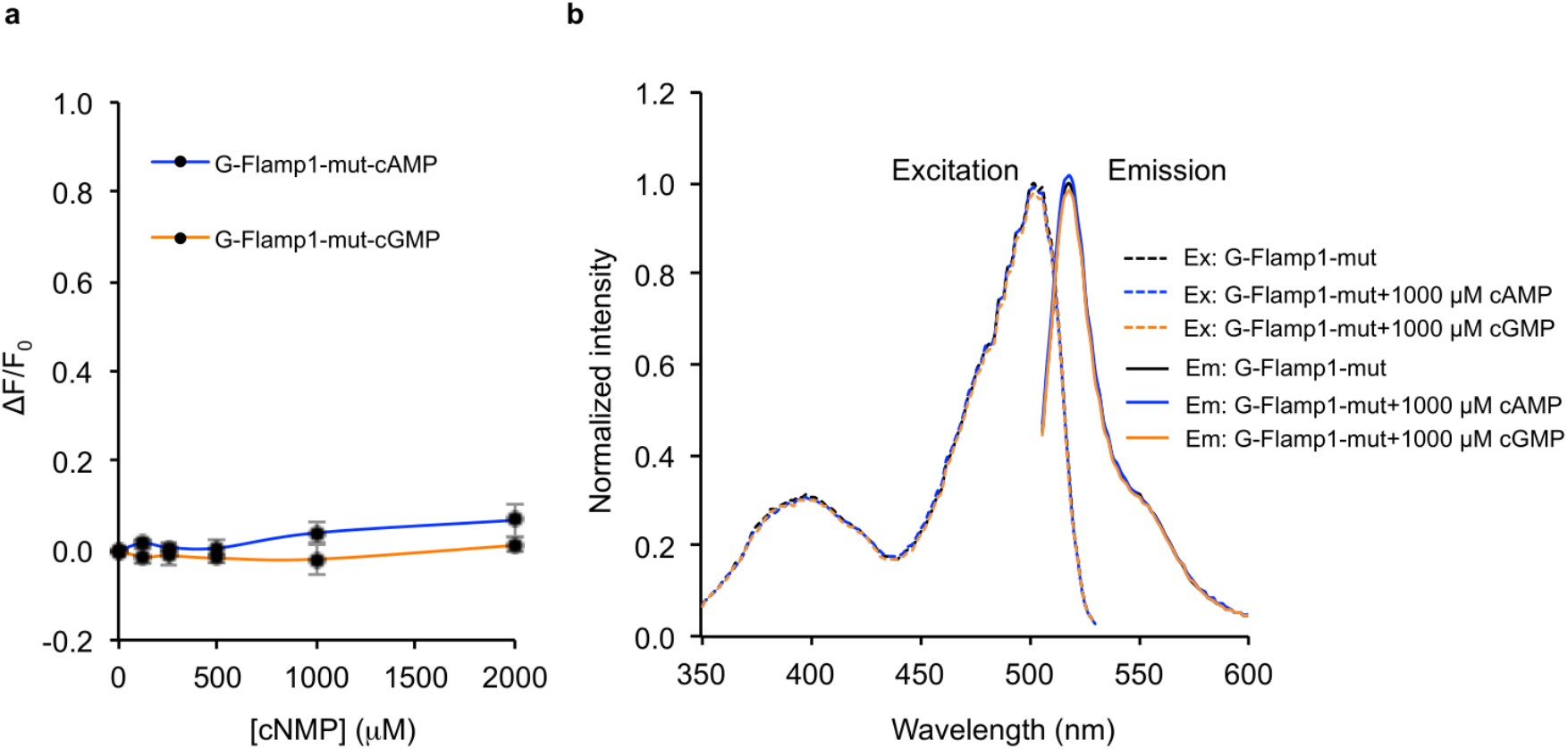
The responses of G-Flamp1-mut to cAMP or cGMP *in vitro*. (a) ΔF/F_0_ of purified G-Flamp1-mut in response to various concentrations of cAMP or cGMP. The fluorescence under excitation at 450 nm was collected. Data are shown as mean ± SEM from 3 independent experiments. (b) Excitation and emission spectra of purified G-Flamp1-mut without cAMP or cGMP (black), with 1000 µM cAMP (blue) and with 1000 µM cGMP (orange). Ex and Em stand for excitation and emission, respectively.

**Supplementary Fig. 14.**
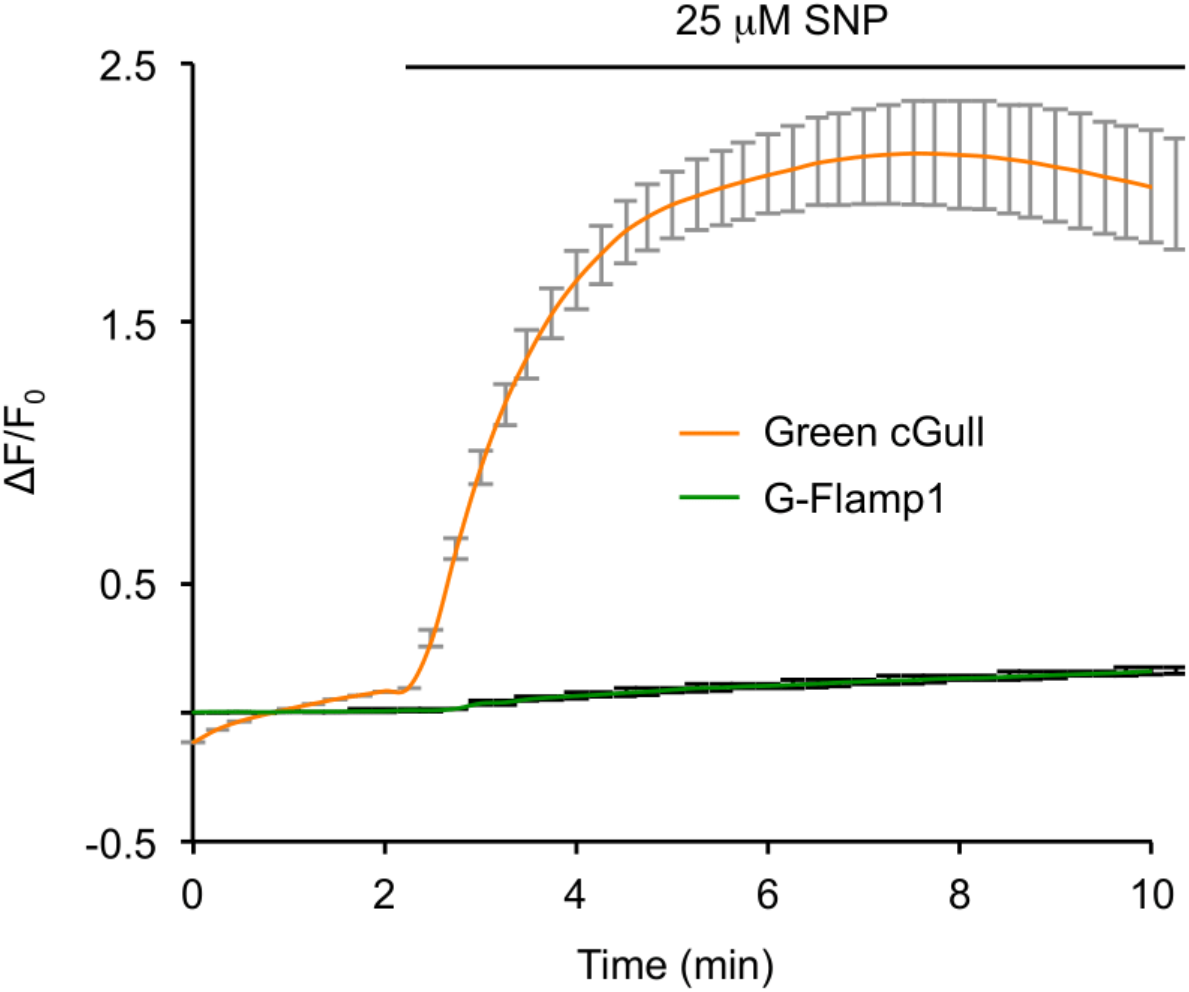
ΔF/F_0_ of Green cGull and G-Flamp1 in response to 25 µM SNP in HEK293T cells. Representative traces of ΔF/F_0_ of HEK293T cells expressing Green cGull or G-Flamp1. Data are shown as mean ± SEM. n = 22 cells for Green cGull and n = 15 cells for G-Flamp1 from 3 cultures for both.

**Supplementary Fig. 15.**
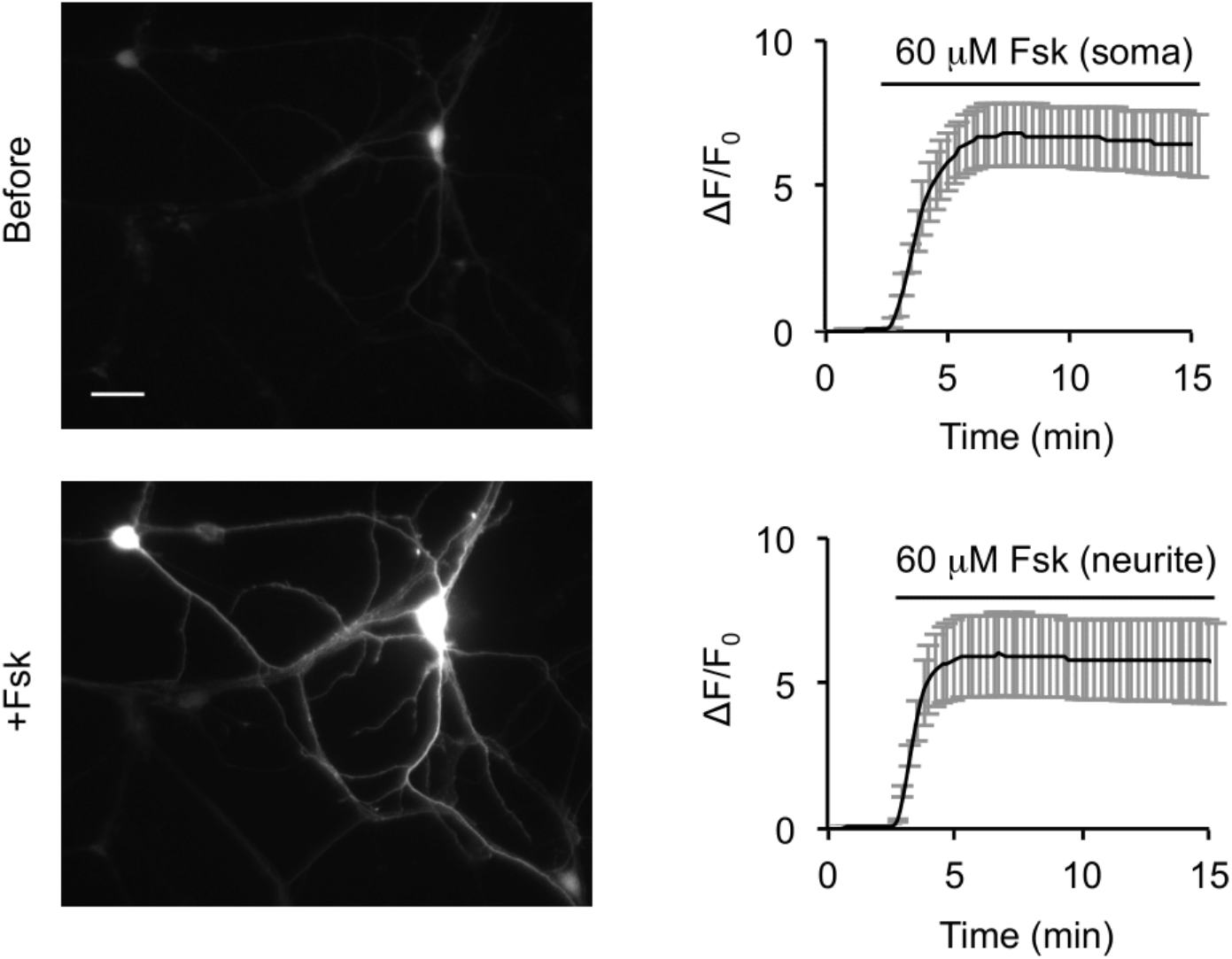
ΔF/F_0_ of G-Flamp1 in cultured cortical neurons in response to 60 µM Fsk. Representative fluorescence images (left) and traces of ΔF/F_0_ (right) of cortical neurons expressing G-Flamp1 in response to 60 µM Fsk. Data are shown as mean ± SEM. n = 6 ROIs of 6 neurons for both soma and neurites. Curves are shown as mean ± SEM. Scale bar: 20 µm.

**Supplementary Fig. 16.**
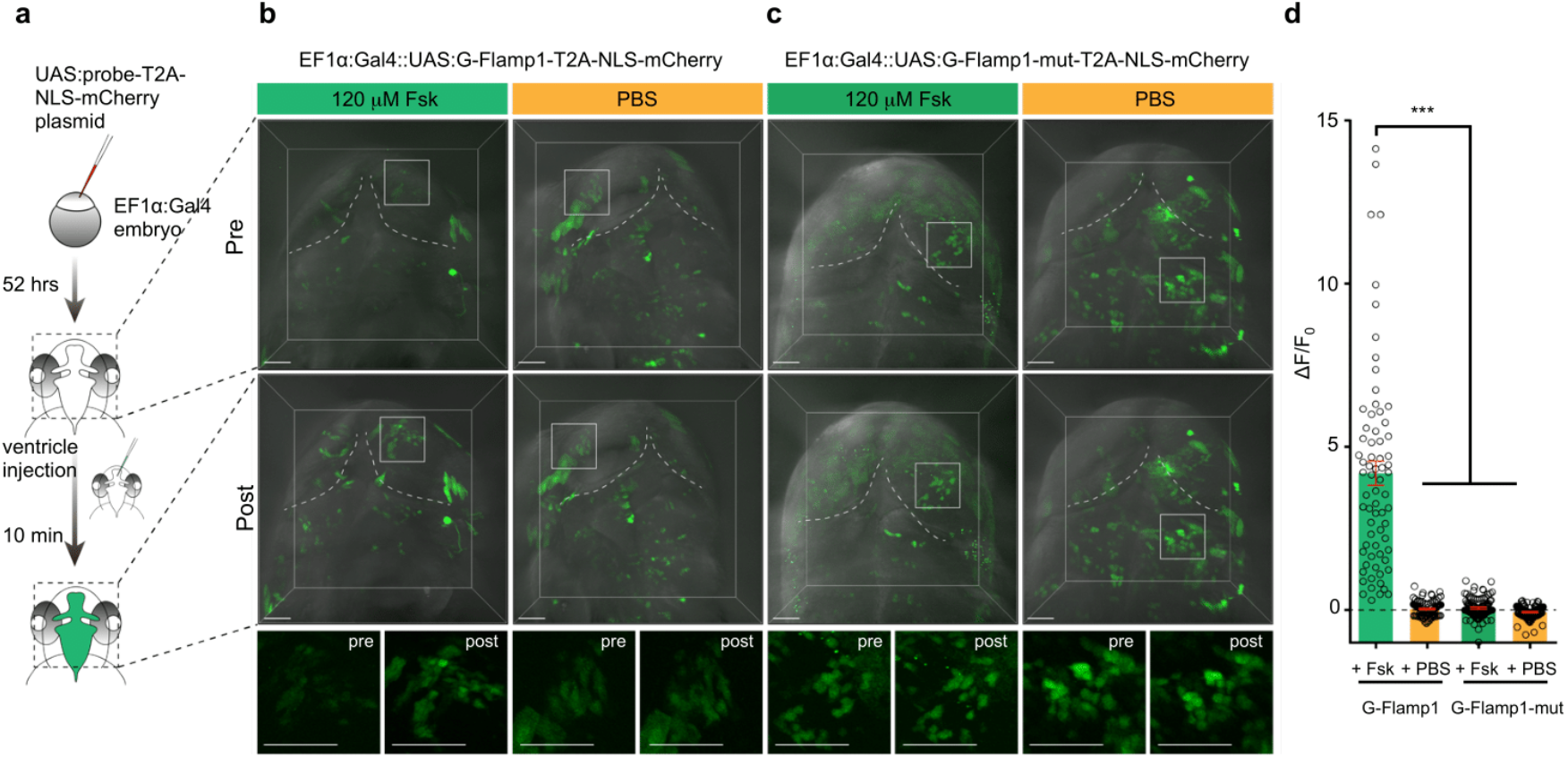
Performance of G-Flamp1 in zebrafish. (a) Schematic drawing for the experiments in zebrafish. (b) Representative fluorescent images of G-Flamp1 before and after 120 µM Fsk or PBS injection. High-magnification images of the boxed areas are shown below. Scale bars: 50 µm. (c) Similar as **b** except that G-Flamp1-mut-T2A-NLS-mCherry plasmid was used. (d) Quantification of ΔF/F_0_ in the above conditions. Data are shown as mean ± SEM overlaid with data points from individual cells. n = 73 cells from 4 animals for G-Flamp1 with Fsk group, 86 cells from 3 animals for G-Flamp1 with PBS group, 93 cells from 3 animals for G-Flamp1-mut with Fsk group, 92 cells from 3 animals for G-Flamp1-mut with PBS group. Two-tailed Student’s *t*-tests were performed. *P* = 1.7 × 10^-13^, 2.1 × 10^-13^ and 7.4 × 10^-14^ between G-Flamp1 with Fsk group and G-Flamp1 with PBS, G-Flamp1-mut with Fsk and G-Flamp1-mut with PBS groups, respectively. \*\*\**P* < 0.001.

### 2. Supplementary Tables

**Supplementary Table 1.** Biophysical and biochemical properties of purified G-Flamp1.

| cAMP<br>sensor | Ex/Em<sup>a</sup><br>(nm),<br>free<sup>b</sup> | Ex/Em<br>(nm),<br>bound<sup>c</sup> | $\Delta F/F_0$<br>(450<br>nm)<sup>d</sup> | $K_d$<br>($\mu M$)<sup>e</sup> | $n_H$<sup>f</sup> | $K_{on}$<br>($\mu M^{-1}s^{-1}$)<sup>g</sup> | $K_{off}$<br>($s^{-1}$)<sup>h</sup> | pKa<sup>i</sup>,<br>free | pKa,<br>bound | EC<sup>j</sup>,<br>free<br>($M^{-1}cm^{-1}$) | EC,<br>bound<br>($M^{-1}cm^{-1}$) | QY<sup>k</sup>,<br>free | QY,<br>bound |
| --- | --- | --- | --- | --- | --- | --- | --- | --- | --- | --- | --- | --- | --- |
| G-Flamp1 | 500/513 | 490/510 | 13.4 | 2.17 | 1.13 | 3.48 | 7.9 | 8.27 | 6.95 | 4374 | 25280 | 0.323 | 0.322 |

<sup>a</sup>Excitation peak/Emission peak. <sup>b</sup>cAMP-free form of G-Flamp1. <sup>c</sup>cAMP-bound form of G-Flamp1. <sup>d</sup>Maximum fluorescence change under 450 nm excitation. <sup>e</sup>Dissociation constant. <sup>f</sup>Hill coefficient. <sup>g</sup>Association rate constant. <sup>h</sup>Dissociation rate constant. <sup>i</sup>The pH at which the fluorescence intensity is half-maximal. <sup>j</sup>Extinction coefficient. <sup>k</sup>Quantum yield.

**Supplementary Table 2.**
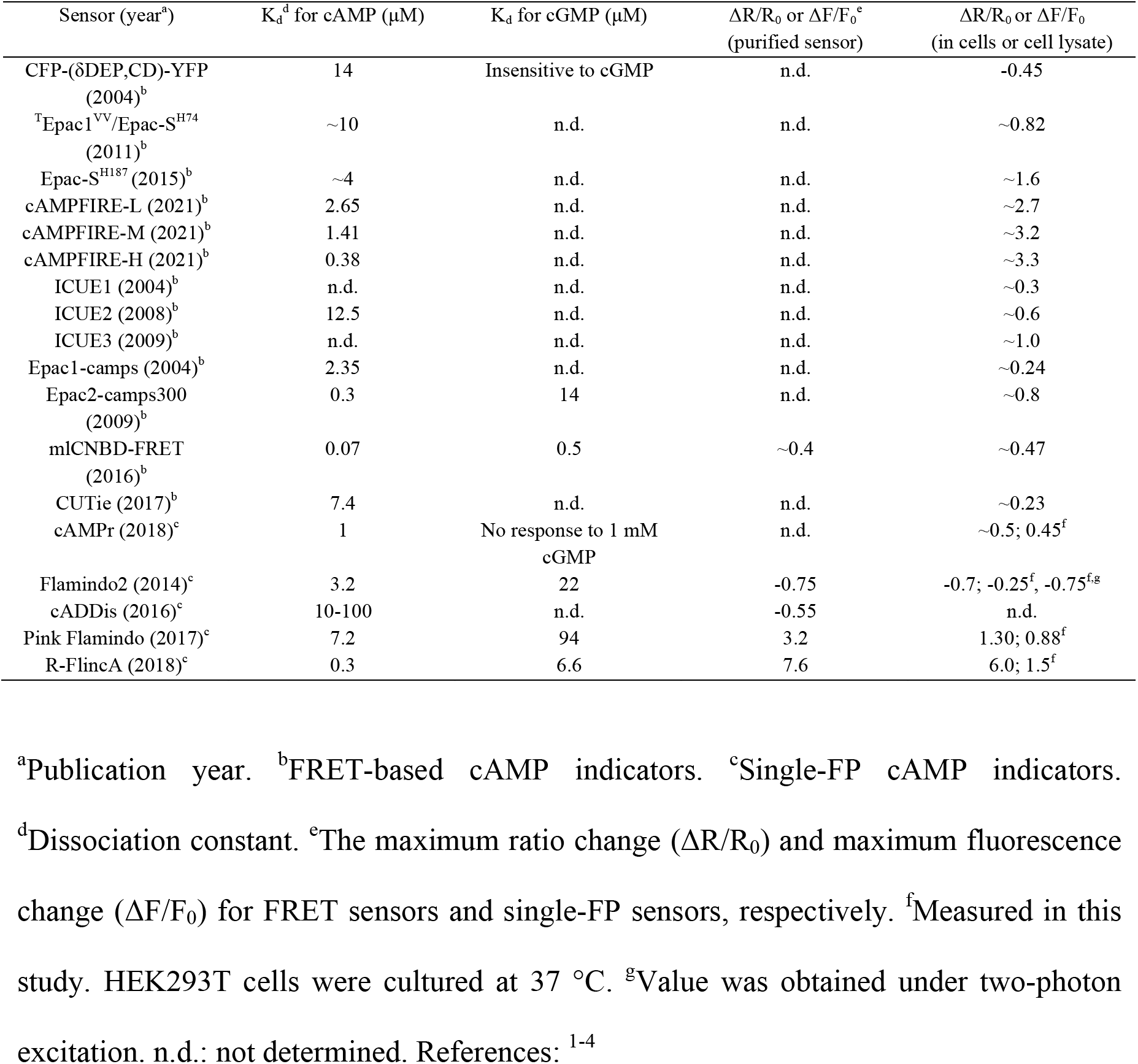
List of current cAMP indicators.

**Supplementary Table 3.** Data collection and structure refinement statistics.

| G-Flamp1 (PDB 6M63) |  |
| --- | --- |
| **Data collection** |  |
| Space group | $P 2_12_12_1$ |
| Cell dimensions |  |
| *a*, *b*, *c* (Å) | 87.84, 94.69, 109.99 |
| $\alpha$, $\beta$, $\gamma$ (°) | 90.00, 90.00, 90.00 |
| Resolution (Å) | 50.00-2.25 (2.29-2.25) * |
| $R_{\text{merge}}^{**}$ | 0.15 (1.01) |
| $I / \sigma I$ | 36.36 (4.45) |
| Completeness (%) | 100.00 (100.00) |
| Redundancy | 14.50 (13.80) |
| **Refinement** |  |
| Resolution (Å) | 47.34-2.25 (2.33-2.25) |
| No. reflections | 43,987 (4,114) |
| $R_{\text{work}}^{\#} / R_{\text{free}}^{\#\#}$ | 0.18/0.22 |
| No. atoms | 5,857 |
| Protein | 5,448 |
| Ligand/ion | 44 |
| Water | 365 |
| *B*-factors | 37.61 |
| Protein | 37.16 |
| Ligand/ion | 28.42 |
| Water | 45.50 |
| R.m.s. deviations |  |
| Bond lengths (Å) | 0.007 |
| Bond angles (°) | 0.900 |
| Ramachandran favored (%) | 98.15 |
| Ramachandran allowed (%) | 1.85 |
| Ramachandran outliers (%) | 0.00 |

\*Statistics for the highest-resolution shell are shown in parentheses.
$R_{\text{merge}}^{**} = \sum_{hkl} \sum_i |I_i(hkl) - \langle I(hkl) \rangle| / \sum_{hkl} \sum_i I_i(hkl)$ , where $I_i(hkl)$ is the intensity measured for the *i* th reflection and $\langle I(hkl) \rangle$ is the average intensity of all reflections with indices *hkl*.
$R_{\text{work}}^{\#} = \sum_{hkl} ||F_{\text{obs}}(hkl) - |F_{\text{calc}}(hkl)|| / \sum_{hkl} |F_{\text{obs}}(hkl)|$ .
$R_{\text{free}}^{\#\#}$ is calculated in an identical manner using 10% of randomly selected reflections that were not included in the refinement.

**Supplementary Table 4.**
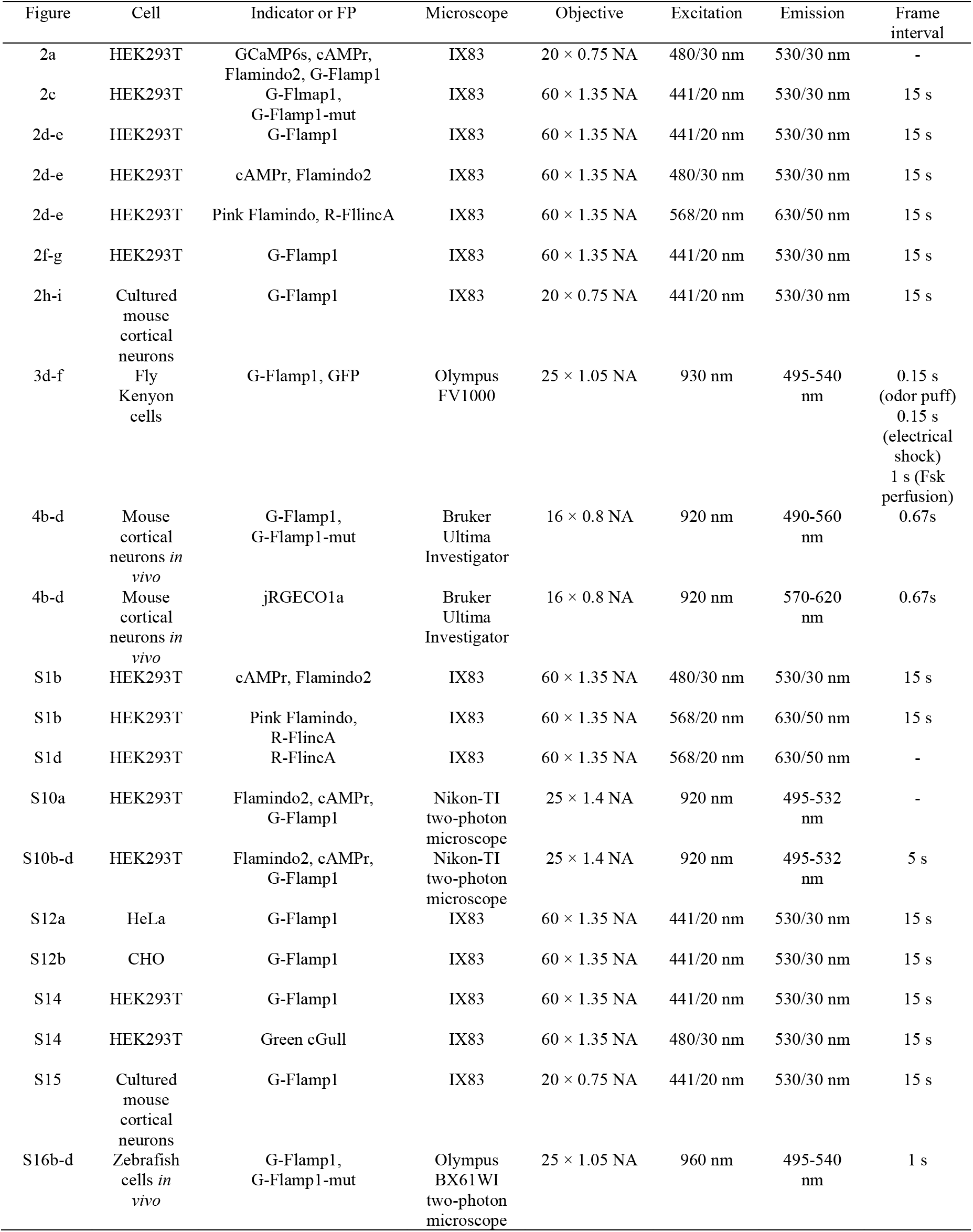
Key parameters for fluorescence imaging data collection.

### 3. Legend for Supplementary Video

#### Supplementary Video 1

Molecular dynamics simulation of cAMP-free G-Flamp1 with a length of 157.50 ns. The yellow residue in the movie is Trp75 and the blue is the chromophore.

## Notes

### Competing Interest Statement

The authors have declared no competing interest.

## References

1. Karpen, J.W. Perspectives on: Cyclic nucleotide microdomains and signaling specificity. J Gen Physiol 143, 5–7 (2014).

2. Sassone-Corsi, P. The cyclic AMP pathway. Cold Spring Harb Perspect Biol 4 (2012).

3. Musheshe, N., Schmidt, M. & Zaccolo, M. cAMP: From Long-Range Second Messenger to Nanodomain Signalling. Trends Pharmacol Sci 39, 209–222 (2018).

4. Surdo, N.C. et al. FRET biosensor uncovers cAMP nano-domains at beta-adrenergic targets that dictate precise tuning of cardiac contractility. Nat Commun 8, 15031 (2017).

5. Bock, A. et al. Optical Mapping of cAMP Signaling at the Nanometer Scale. Cell 182, 1519–1530.e1517 (2020).

6. Lin, M.Z. & Schnitzer, M.J. Genetically encoded indicators of neuronal activity. Nature Neuroscience 19, 1142–1153 (2016).

7. Zhang, Q. et al. Designing a Green Fluorogenic Protease Reporter by Flipping a Beta Strand of GFP for Imaging Apoptosis in Animals. J Am Chem Soc 141, 4526–4530 (2019).

8. Liu, W. et al. Genetically encoded single circularly permuted fluorescent protein-based intensity indicators. Journal of Physics D: Applied Physics 53, 113001 (2020).

9. Marvin, J.S., Schreiter, E.R., Echevarria, I.M. & Looger, L.L. A genetically encoded, high-signal-to-noise maltose sensor. Proteins 79, 3025–3036 (2011).

10. Marvin, J.S. et al. An optimized fluorescent probe for visualizing glutamate neurotransmission. Nat Methods 10, 162–170 (2013).

11. Odaka, H., Arai, S., Inoue, T. & Kitaguchi, T. Genetically-encoded yellow fluorescent cAMP indicator with an expanded dynamic range for dual-color imaging. PLoS One 9, e100252 (2014).

12. Christopher R. Hackley, E.O.M., Justin Blau cAMPr: A single-wavelength fluorescent sensor for cyclic AMP. SCIENCE SIGNALING 11, eaah3738 (2018).

13. Harada, K. et al. Red fluorescent protein-based cAMP indicator applicable to optogenetics and in vivo imaging. Sci Rep 7, 7351 (2017).

14. Ohta, Y., Furuta, T., Nagai, T. & Horikawa, K. Red fluorescent cAMP indicator with increased affinity and expanded dynamic range. Sci Rep 8, 1866 (2018).

15. Nimigean, C.M., Shane, T. & Miller, C. A cyclic nucleotide modulated prokaryotic K+ channel. J Gen Physiol 124, 203–210 (2004).

16. Mukherjee, S. et al. A novel biosensor to study cAMP dynamics in cilia and flagella. Elife 5 (2016).

17. Chen, T.-W. et al. Ultrasensitive fluorescent proteins for imaging neuronal activity. Nature 499, 295–300 (2013).

18. Pedelacq, J.D., Cabantous, S., Tran, T., Terwilliger, T.C. & Waldo, G.S. Engineering and characterization of a superfolder green fluorescent protein. Nat Biotechnol 24, 79–88 (2006).

19. Griesbeck, O., Baird, G.S., Campbell, R.E., Zacharias, D.A. & Tsien, R.Y. Reducing the Environmental Sensitivity of Yellow Fluorescent Protein. Journal of Biological Chemistry 276, 29188–29194 (2001).

20. Altieri, S.L. et al. Structural and Energetic Analysis of Activation by a Cyclic Nucleotide Binding Domain. Journal of Molecular Biology 381, 655–669 (2008).

21. Permyakov, E.A., Barnett, L.M., Hughes, T.E. & Drobizhev, M. Deciphering the molecular mechanism responsible for GCaMP6m’s Ca2+-dependent change in fluorescence. Plos One 12, e0170934 (2017).

22. Özel, R.E., Lohith, A., Mak, W.H. & Pourmand, N. Single-cell intracellular nano-pH probes. RSC Advances 5, 52436–52443 (2015).

23. Borner, S. et al. FRET measurements of intracellular cAMP concentrations and cAMP analog permeability in intact cells. Nat Protoc 6, 427–438 (2011).

24. Jiang, J.Y., Falcone, J.L., Curci, S. & Hofer, A.M. Interrogating cyclic AMP signaling using optical approaches. Cell Calcium 64, 47–56 (2017).

25. Campbell, R.E. et al. A monomeric red fluorescent protein. Proc Natl Acad Sci U S A 99, 7877–7882 (2002).

26. Shen, Y. et al. A genetically encoded Ca(2+) indicator based on circularly permutated sea anemone red fluorescent protein eqFP578. BMC Biol 16, 9 (2018).

27. Sabatini, B.L. & Tian, L. Imaging Neurotransmitter and Neuromodulator Dynamics In Vivo with Genetically Encoded Indicators. Neuron 108, 17–32 (2020).

28. Marvin, J.S. et al. A genetically encoded fluorescent sensor for in vivo imaging of GABA. Nature Methods 16, 763–770 (2019).

29. Laviv, T. et al. In Vivo Imaging of the Coupling between Neuronal and CREB Activity in the Mouse Brain. Neuron 105, 799–812.e795 (2020).

30. Chao Qi, S.S., Ohad Medalia, Volodymyr M. Korkhov The structure of a membrane adenylyl cyclase bound to an activated stimulatory G protein. Science 364, 389–394 (2019).

31. Russwurm, M. & Koesling, D. Measurement of cGMP-generating and -degrading activities and cGMP levels in cells and tissues: Focus on FRET-based cGMP indicators. Nitric Oxide 77, 44–52 (2018).

32. Matsuda, S. et al. Generation of a cGMP Indicator with an Expanded Dynamic Range by Optimization of Amino Acid Linkers between a Fluorescent Protein and PDE5alpha. ACS Sens 2, 46–51 (2017).

33. Kandel, E.R. The molecular biology of memory storage: a dialogue between genes and synapses. Science 294, 1030–1038 (2001).

34. Heisenberg, M. Mushroom body memoir: from maps to models. Nat Rev Neurosci 4, 266–275 (2003).

35. Livingstone, M.S., Sziber, P.P. & Quinn, W.G. Loss of calcium/calmodulin responsiveness in adenylate cyclase of rutabaga, a Drosophila learning mutant. Cell 37, 205–215 (1984).

36. Zars, T., Wolf, R., Davis, R. & Heisenberg, M. Tissue-specific expression of a type I adenylyl cyclase rescues the rutabaga mutant memory defect: in search of the engram. Learn Mem 7, 18–31 (2000).

37. Ma, L. et al. A Highly Sensitive A-Kinase Activity Reporter for Imaging Neuromodulatory Events in Awake Mice. Neuron 99, 665–679.e665 (2018).

38. Lee, S.J. et al. Cell-type-specific asynchronous modulation of PKA by dopamine in learning. Nature 590, 451–456 (2020).

39. Schultz, W., Dayan, P. & Montague, P.R. A neural substrate of prediction and reward. Science 275, 1593–1599 (1997).

40. Sun, F. et al. A Genetically Encoded Fluorescent Sensor Enables Rapid and Specific Detection of Dopamine in Flies, Fish, and Mice. Cell 174, 481–496.e419 (2018).

41. Oe, Y. et al. Distinct temporal integration of noradrenaline signaling by astrocytic second messengers during vigilance. Nature Communications 11 (2020).

42. Zhang, S.X. et al. Hypothalamic dopamine neurons motivate mating through persistent cAMP signalling. Nature 597, 245–249 (2021).

43. Massengill, C.I., et al. Highly sensitive genetically-encoded sensors for population and subcellular imaging of cAMP in vivo. *Preprint at* bioRxiv https://www.biorxiv.org/content/10.1101/2021.08.27.457999v1 (2021).

44. Zong, W. et al. Miniature two-photon microscopy for enlarged field-of-view, multi-plane and long-term brain imaging. Nature Methods 18, 46–49 (2021).

45. Muntean, B.S. et al. Interrogating the Spatiotemporal Landscape of Neuromodulatory GPCR Signaling by Real-Time Imaging of cAMP in Intact Neurons and Circuits. Cell Rep 24, 1081–1084 (2018).

46. Plattner, F. et al. The role of ventral striatal cAMP signaling in stress-induced behaviors. Nature Neuroscience 18, 1094–1100 (2015).

47. Nasu, Y., Shen, Y., Kramer, L. & Campbell, R.E. Structure-and mechanism-guided design of single fluorescent protein-based biosensors. Nature Chemical Biology 17, 509–518 (2021).

48. Zhao, Y. et al. An Expanded Palette of Genetically Encoded Ca2+ Indicators. Science 333, 1888–1891 (2011).

49. Qian, Y. et al. A genetically encoded near-infrared fluorescent calcium ion indicator. Nat Methods 16, 171–174 (2019).

50. Fosque, B.F. et al. Labeling of active neural circuits in vivo with designed calcium integrators. Science 347, 755–760 (2015).

51. Dana, H. et al. Sensitive red protein calcium indicators for imaging neural activity. Elife 5 (2016).

52. Chu, J. et al. A bright cyan-excitable orange fluorescent protein facilitates dual-emission microscopy and enhances bioluminescence imaging in vivo. Nat Biotechnol 34, 760–767 (2016).

53. Tian, L. et al. Imaging neural activity in worms, flies and mice with improved GCaMP calcium indicators. Nature Methods 6, 875–881 (2009).

54. Datsenko, K.A. & Wanner, B.L. One-step inactivation of chromosomal genes in Escherichia coli K-12 using PCR products. Proc Natl Acad Sci U S A 97, 6640–6645 (2000).

55. Shinoda, H. et al. Acid-Tolerant Monomeric GFP from Olindias formosa. Cell Chemical Biology 25, 330–338.e337 (2018).

56. Slepenkov, S.V., Darzynkiewicz, E. & Rhoads, R.E. Stopped-flow kinetic analysis of eIF4E and phosphorylated eIF4E binding to cap analogs and capped oligoribonucleotides: evidence for a one-step binding mechanism. J Biol Chem 281, 14927–14938 (2006).

57. Nausch, L.W., Ledoux, J., Bonev, A.D., Nelson, M.T. & Dostmann, W.R. Differential patterning of cGMP in vascular smooth muscle cells revealed by single GFP-linked biosensors. Proc Natl Acad Sci U S A 105, 365–370 (2008).

58. Minor, W., Cymborowski, M., Otwinowski, Z. & Chruszcz, M. HKL-3000: the integration of data reduction and structure solution--from diffraction images to an initial model in minutes. Acta Crystallogr D Biol Crystallogr 62, 859–866 (2006).

59. McCoy, A.J., et al. Phaser crystallographic software. J Appl Crystallogr 40, 658–674 (2007).

60. Adams, P.D. et al. PHENIX: a comprehensive Python-based system for macromolecular structure solution. Acta Crystallogr D Biol Crystallogr 66, 213–221 (2010).

61. Emsley, P., Lohkamp, B., Scott, W.G. & Cowtan, K. Features and development of Coot. Acta Crystallogr D Biol Crystallogr 66, 486–501 (2010).

62. Krieger, E. & Vriend, G. New ways to boost molecular dynamics simulations. J Comput Chem 36, 996–1007 (2015).

63. Beaudoin, G.M.J. et al. Culturing pyramidal neurons from the early postnatal mouse hippocampus and cortex. Nature Protocols 7, 1741–1754 (2012).

64. Chu, J. et al. Non-invasive intravital imaging of cellular differentiation with a bright red-excitable fluorescent protein. Nature Methods 11, 572–578 (2014).

65. Cantu, D.A. et al. EZcalcium: Open-Source Toolbox for Analysis of Calcium Imaging Data. Frontiers in Neural Circuits 14 (2020).

## References

1. Jiang, J.Y., Falcone, J.L., Curci, S. & Hofer, A.M. Interrogating cyclic AMP signaling using optical approaches. Cell Calcium 64, 47–56 (2017).

2. Klausen, C., Kaiser, F., Stüven, B., Hansen, J.N. & Wachten, D. Elucidating cyclic AMP signaling in subcellular domains with optogenetic tools and fluorescent biosensors. Biochemical Society Transactions 47, 1733–1747 (2019).

3. Dikolayev, V., Tuganbekov, T. & Nikolaev, V.O. Visualizing Cyclic Adenosine Monophosphate in Cardiac Microdomains Involved in Ion Homeostasis. Front Physiol 10, 1406 (2019).

4. Massengill, C.I., et al. Highly sensitive genetically-encoded sensors for population and subcellular imaging of cAMP in vivo. *Preprint at* bioRxiv https://www.biorxiv.org/content/10.1101/2021.08.27.457999v1 (2021).

